# Retron-Eco9 ADP-ribosylates release factors to disrupt translation during phage infection

**DOI:** 10.64898/2026.11.25.769155

**Authors:** Christina R. Beck, Mira B. May, Joseph H. Davis, Michael T. Laub

## Abstract

Retrons are a major form of anti-phage defense in bacteria, but how they provide defense remains incompletely understood. The most common retron effector is a predicted nucleoside 2’-deoxyribosyltransferase (NDT). Here, we demonstrated that the NDT of retron-Eco9 is, in fact, a protein histidine ADP-ribosyltransferase that associates with the bacterial ribosome through an interaction with the stalk proteins bL7/bL12. Through this interaction, the effector ADP-ribosylates translation release factors RF1 and RF2 adjacent to the conserved GGQ motifs that catalyze peptidyl-tRNA ester bond hydrolysis, resulting in increased stop codon readthrough and frameshifting. Using ribosome profiling, cryoEM, and *in vitro* reconstitution assays, we demonstrate that the modification of RF1 and RF2 severely disrupts translation in cells, leading to collided ribosomes at the 3’ ends of transcripts, aberrant termination, and increased ribosome rescue. These disruptions to translation ultimately prevent phage replication and abort an infection. Although many defense systems block translation by degrading tRNA or mRNA, our results now demonstrate how post-translational modification of translation factors can also provide potent defense, while also revealing histidine ADP-ribosyltransferase activity as the biochemical basis of the largest class of retrons.

## INTRODUCTION

Shutting off translation is a common and effective strategy for aborting viral infection of host cells throughout biology. For instance, some viruses that infect mammalian cells activate protein kinase R, which phosphorylates eIF2a to block translation^1,2^, while others stimulate oligoadenylate synthetase, which, by driving activation of RNase L, leads to degradation of viral and cellular mRNAs and thereby blocks translation^1,3,4^. In bacteria, many anti-phage defense systems also inhibit translation to restrict viral replication^5–10^. In most cases, this inhibition arises from effector proteins that target or degrade tRNA or mRNA rather than targeting ribosomal proteins or translation factors. The Lit protease, which provides defense against T4 phage, cleaves EF-Tu to block translation and abort infection^11^, but whether other components of the translation machinery are targeted by bacterial defense systems is largely unknown.

Retrons are an abundant class of anti-phage defense systems, canonically comprised of a non-coding RNA (ncRNA), a reverse transcriptase (RT), and an effector protein^12,13^. The RT typically synthesizes DNA, called multi-copy single-stranded DNA (msDNA), through a covalent linkage to the ncRNA template. The msDNA, together with the RT, binds and restrains the effector. Phage infection can disrupt this complex, leading to release of the effector. Retron-encoded effectors exhibit diverse catalytic activities across different retron classes, including tRNases and NADases^12,14–16^. However, the most common effector, found in type II retrons, is a predicted nucleoside 2’-deoxyribosyltransferase (NDT)^12^. Despite its prevalence, the target and activity of many effectors within this class are unknown.

Here, we characterized Retron-Eco9 and show that the effector rapidly inhibits translation, but not by targeting tRNAs or mRNAs. Instead, this effector ADP-ribosylates the release factors RF1 and RF2 to interfere with translation termination, with evidence pointing to both a failure to properly terminate translation and additional readthrough, often with frameshifting. Further, the disruption of termination induces ribosome collisions and ribosome accumulation at the ends of mRNAs, which, in turn, triggers ribosome rescue pathways, including the SmpB-*ssrA* pathway. Our work highlights the first defense protein known to target translation release factors, and reveals how it prompts major translation dysfunction prior to cell death.

## RESULTS

### Effector-Eco9 defends against phage by shutting off translation

We focused on Retron-Eco9, a member of the type-II family of retrons, that was first identified in the *Escherichia coli* isolate NILS-16 through homology to the related system Retron-Sen2^12^ (**Figure 1A**). Retron-Eco9 features an effector (previously called RcaT but hereafter effector-Eco9) that contains a winged helix-turn-helix (HTH) domain and a predicted nucleoside 2’-deoxyribosyltransferase (NDT)-like domain, the most common effector domain in retrons^13^ (**Figure S1A-B**). To measure anti-phage defense provided by Retron-Eco9, we expressed it from its native promoter on a low-copy plasmid in *E. coli* MG1655 and then challenged the strain with 12 diverse coliphages. Retron-Eco9 decreased the efficiency of plaquing (EOP) for T5, T7, SECFΦ17, λ_vir_, and Lust by 10-to 10^5^-fold (**Figure 1B, S1C**), indicating the system provides strong defense against a wide range of phages. Substituting the predicted catalytic glutamate, E108, in effector-Eco9 with alanine restored plaquing by Lust (**Figure 1C, S1D**) indicating that activity of the NDT-like domain is necessary for defense.

**Figure 1.**
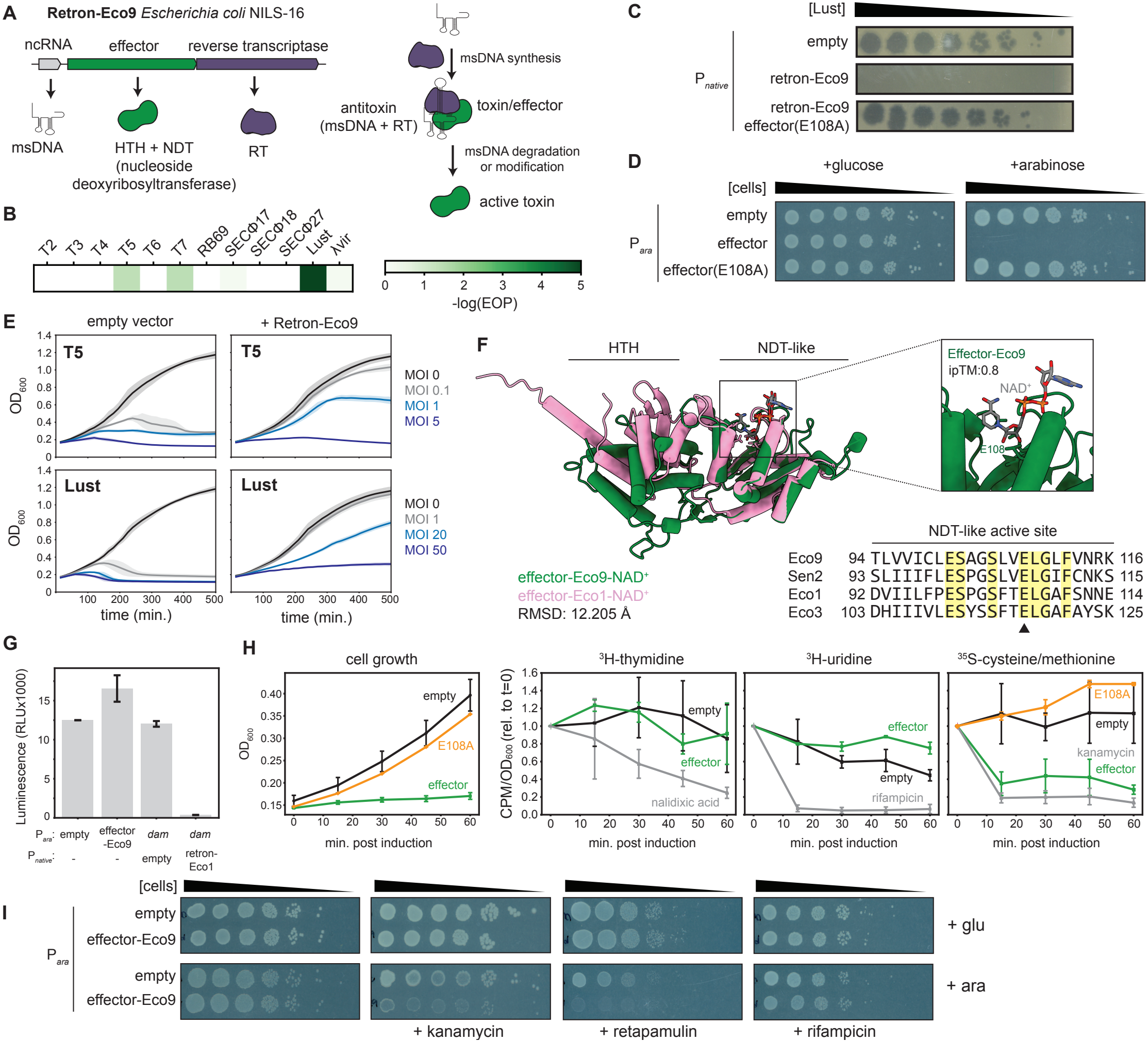
Effector-Eco9 defends against phage by disrupting translation. **(A)** Schematic of Retron-Eco9 operon and mechanism of toxin activation. **(B)** Heatmap depicting efficiency of plaquing (EOP) for the indicated phages when infecting cells producing Retron-Eco9. **(C)** Plaquing of serial dilutions of Lust phage on lawns of cells containing plasmids expressing Retron-Eco9, Retron-Eco9 (Effector E108A), or an empty vector. **(D)** Cell viability assessed by plating of serial dilutions of strains expressing effector-Eco9 or the effector-Eco9(E108A) mutant from an arabinose-inducible promoter. **(E)** Growth of cells producing Retron-Eco9 or an empty vector following infection with T5 or Lust at the indicated MOIs. Data are mean ± s.d. (shaded region) of four total measurements from two biological replicates. **(F)** left, overlay of AF3-predicted effector-Eco9-NAD^+^ complex and effector-Eco1-NAD^+^ complex; top right inset, NAD^+^ bound to the effector-Eco9 active site predicted by AF3; bottom right, alignment of active site residues in closely related type II retron effectors (yellow highlight: conserved residues, black arrowhead: catalytic glutamate). **(G)** NADase activity assessed by a luminescence-based assay in cells expressing effector-Eco9, *E. coli dam*, Retron-Eco1 and *E. coli dam*, or an empty vector. Data are mean ± s.d. of six total replicates from two biological samples. **(H)** left, OD_600_ of cells expressing effector-Eco9 or effector-Eco9(E108A) from an arabinose-inducible promoter at the times indicated post-addition of arabinose; right, cells were concurrently pulse-labeled with ^3^H-thymidine, ^3^H-uridine, or ^35^S-cysteine/methionine mix at the indicated times. Nalidixic acid, rifampicin, and kanamycin treated cells were included as controls. CPM values at each timepoint were normalized by OD_600_ and are presented relative to t=0. Data are mean ± s.d. of two biological replicates. **(I)** Cell viability assessed by plating serial dilutions of strains expressing effector-Eco9 from an abinose-inducible promoter in the presence of sub-lethal concentrations of arabinose and kanamycin, retapamulin, or rifampicin (ara: arabinose, glu: glucose).

To assess whether Retron-Eco9 behaves like a toxin-antitoxin (TA) system, we induced effector-Eco9 from an arabinose-inducible promoter on a plasmid in wild-type *E. coli*. The effector caused a nearly complete loss of colony forming units, consistent with previous reports^13^ (**Figure 1D**), in a manner dependent on the predicted catalytic residue E108. Thus, effector-Eco9 is toxic when expressed alone without the RT and msDNA components that normally restrain its activity, as in canonical TA systems.

To confirm that Retron-Eco9 functions through an abortive infection mechanism^17^, we infected cells harboring Retron-Eco9 with T5 or Lust at a range of multiplicities of infections (MOIs). We observed defense only at low MOIs, consistent with Retron-Eco9 functioning through an abortive infection mechanism (**Figure 1E**) in which infected cells do not survive but prevent production of mature virions, thereby sparing uninfected cells in the population.

The Retron-Eco1 effector functions as an NADase and possibly as a DNA ADP-ribosyltransferase^14^. Effector-Eco9 has strong sequence similarity to the effectors of retrons Eco1, Eco3, and Sen2^13^, and AlphaFold3^18^ produced a high-confidence prediction for effector-Eco9 bound to NAD^+^ in both monomeric and dimeric states (ipTM: 0.8 (monomer), 0.89-0.90 (dimer), **Figure 1F, S1A-B**). To directly test whether effector-Eco9 has NADase activity, we measured NAD^+^ levels through a luminescence-based assay in cells overproducing effector-Eco9 (**Figure 1G**). We observed no significant decrease in luminescence, whereas activating effector-Eco1 by co-producing *E. coli* Dam (DNA adenine methyltransferase)^19^ led to a decrease in luminescence > 10,000-fold. We conclude that effector-Eco9 is not an NADase and instead may use NAD^+^ for ADP-ribosylation.

To determine what cellular process effector-Eco9 disrupts, we performed pulse-labeling assays to monitor DNA replication, transcription, and translation (**Figure 1H**). Following induction of effector-Eco9 from an arabinose-inducible promoter, we measured cell growth (OD_600_) and the incorporation of ^3^H-thymidine, ^3^H-uridine, or ^35^S-cysteine/ methionine to assess DNA replication, transcription, and translation, respectively. Upon induction, effector-Eco9 almost immediately halted cell growth and led to a dramatic decrease in ^35^S-cysteine/methionine incorporation, comparable to that observed when treating control cells with kanamycin, an antibiotic that targets the bacterial ribosome.

For ^3^H-uridine and ^3^H-thymidine incorporation, we observed no substantial differences between cells producing effector-Eco9 or those harboring an empty vector control, whereas cells treated with rifampicin or nalidixic acid, which are transcription and DNA replication inhibitors, respectively, blocked label incorporation, as expected.

If effector-Eco9 targets translation, we reasoned that it would be synthetically lethal with sub-MIC levels of ribosome-targeting antibiotics. Indeed, in combination with sub-MIC levels of kanamycin or retapamulin, sublethal levels of effector-Eco9 were substantially more toxic, as evident by extremely small colonies (**Figure 1I**). In contrast, sub-MIC levels of rifampicin, which targets transcription, did not induce the same reduction in colony sizes. Collectively, our results indicate that effector-Eco9 causes cellular toxicity by disrupting translation.

### Effector-Eco9 ADP-ribosylates RF1 and RF2

To test whether effector-Eco9 alone is sufficient to block translation, we purified effector-Eco9-3XFLAG from yeast cells, where overexpression is not toxic. Purified effector-Eco9, but not a E108A variant predicted to be catalytically inactive, strongly inhibited production of the reporter protein DHFR in an *in vitro* transcription/translation (IVTT) reaction, with inhibition dependent on NAD^+^ (**Figure 2A**).

**Figure 2.**
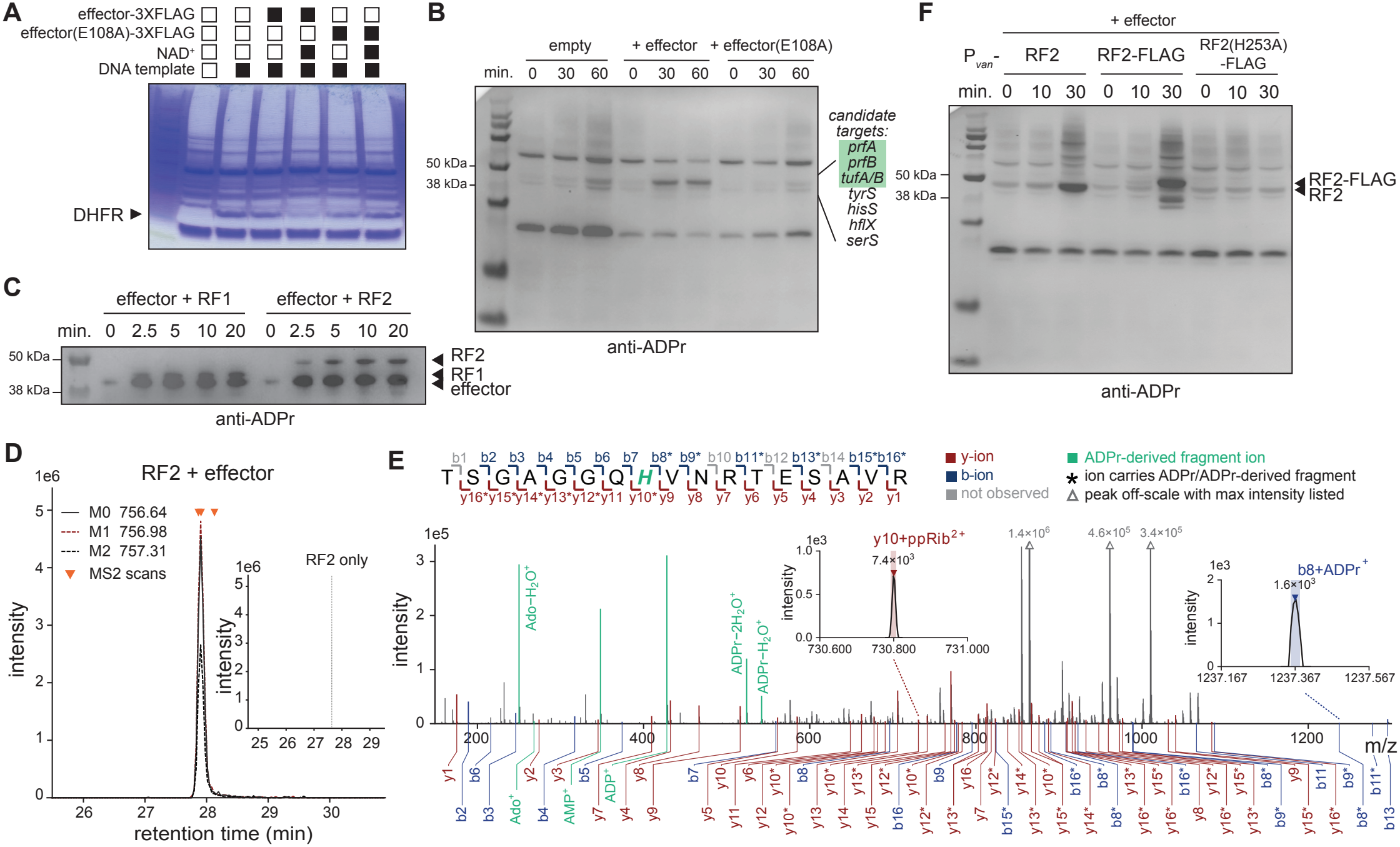
Effector-Eco9 ADP-ribosylates RF1 and RF2. **(A)** *In vitro* transcription-translation (IVTT) assays using DHFR production from a DNA template as the readout of expression activity. Purified Effector-Eco9, Effector-Eco9(E108A), and NAD^+^ were added to the reaction where indicated. Representative of two biological replicates. **(B)** Immunoblot of ADP-ribosylated proteins from cell lysates containing plasmids expressing effector-Eco9 or the E108A mutant from an arabinose-inducible promoter at the times indicated post-arabinose addition. Right, candidate targets within the 38-50 kDa size range. **(C)** Anti-ADPr immunoblot of *in vitro* reactions containing purified effector, RF1 or RF2, and NAD^+^. Proteins were incubated for the indicated times before the reaction was quenched. **(D)** Extracted ion chromatograms of the M0, M1, and M2 isotopomers for the ADP-ribosylated TSGAGGQHVNRTESAVR^3+^ precursor from the RF2 + effector sample and the retention-time adjusted window for the RF2 only sample (inset). Orange triangles mark DDA-triggered MS2 scans. **(E)** Fragment ladder of the TSGAGGQHVNRTESAVR peptide (RF2 residues 246-262) with b and y-ions indicated with brackets (top). Annotated fragmentation spectra (MS2) with y- and b-ions bearing modifications common in HCD-based fragmentation of ADPr noted with asterisk. Released ADPr marker ions (green) noted, with water loss labeled ‘-H_2_O’. Insets depict spectra for b8 and y10-ions bearing ADPr-derived modifications, which jointly localize the modification to H253. Colored rectangle marks 10 ppm mass window. **(F)** Immunoblots of ADP-ribosylated proteins from cell lysates containing plasmids expressing effector-Eco9 from an arabinose-inducible promoter and RF2, RF2-FLAG, or RF2(H253A)-FLAG from a vanillic acid-inducible promoter. Cells were harvested at the indicated times post-arabinose addition.

To probe tRNA as a potential target, tRNAs purified from cells producing effector-Eco9 were added to an IVTT reaction lacking tRNAs. We found that the tRNAs from cells producing effector-Eco9 or an empty vector were both functional, as evidenced by the robust accumulation of DHFR-FLAG (**Figure S2A**). In contrast, tRNAs from cells producing the toxic domain of CapRel^SJ46^, which pyrophosphorylates the 3’ end of tRNAs to prevent tRNA charging^20^, prevented DHFR production. Thus, effector-Eco9 likely does not target tRNAs.

Although NDT enzymes canonically transfer deoxyribosyl moieties between nucleobases, we hypothesized that effector-Eco9 instead transfers ADP-ribose to a ribosomal RNA or protein involved in translation. To probe for ADP-ribosylation, we harvested lysates from cells producing effector-Eco9 and performed a dot blot on total RNA and immunoblotting on proteins using an ADP-ribose-specific antibody. While there was no signal on the dot blot (**Figure S2B**), immunoblotting of proteins revealed one new band in cells producing effector that was absent in empty vector and E108A mutant controls (**Figure 2B**). In the approximate molecular weight range of this ADP-ribosylated band, there are three essential translation factors: PrfA (RF1), PrfB (RF2), and TufA/B (EF-Tu). To assess whether these factors are direct targets of effector-Eco9, we first mixed purified effector with NAD^+^ and purified RF1 or RF2. We then used immunoblots with the ADP-ribose-specific antibody, observing robust modification of both RF1 and RF2, as well as the effector itself (**Figure 2C**). To consider EF-Tu as a target, we expressed *tufA*-FLAG from a constitutive promoter on a high-copy plasmid in a Δ*tufA*Δ*tufB* strain also engineered to produce effector from an arabinose-inducible promoter; immunoblotting lysates from these cells did not reveal any detectable ADP ribosylation of TufA-FLAG (**Figure S2C**).

To determine the specific residue that effector-Eco9 ADP ribosylates on RF1 and RF2, we performed mass spectrometry on *in vitro* reactions containing purified effector, RF2, and NAD^+^ (**Figure 2D-E**). This analysis revealed that H253 was ADP-ribosylated, in an effector-dependent manner. Notably, H253 is directly adjacent to the ^250^GGQ^252^ motif that normally extends into the peptidyl transferase center (PTC) and catalyzes peptidyl-tRNA hydrolysis (**Figure S2D**)^21^, suggesting that ADP-ribosylation of H253 prevents translational termination by RF2 (**Figure S2D**).

To establish that effector-Eco9 ADP-ribosylates RF2 on H253 *in vivo*, we expressed *prfB*-3XFLAG, *prfB(H253A)*-3XFLAG, or wild-type *prfB* in cells also expressing the effector from a plasmid under an arabinose-inducible promoter. Using immunoblots to detect ADP ribosylation, we found that wild-type RF2 was modified 30 minutes post-effector induction (**Figure 2F, S2E-F**). RF2-3XFLAG was also robustly ADP-ribosylated after 30 minutes, with the band shifted upward roughly 3 kDa, whereas cells producing the H253A variant no longer produced a detectable ADPr signal. We conclude that RF2 is ADP-ribosylated *in vivo* by effector-Eco9 on H253. We repeated the same immunoblots but for cells expressing *prfA*-3XFLAG. Following effector induction, we detected ADP-ribosylation of wild-type RF1 and RF1-3XFLAG, but at levels much lower than with RF2-3XFLAG (**Figure S2F**).

### ADP-ribosylation of RF1 and RF2 disrupts translation

To determine if ADP-ribosylation of RF1 or RF2 affects translation, we added purified effector to *in vitro* transcription-translation (IVTT) reactions and monitored production of an HA-DHFR reporter. Importantly, the DNA template encoding HA-DHFR was engineered to use one of three stop codons: TAA, recognized by RF1 and RF2; TAG, recognized by RF1; or TGA, recognized by RF2. Strikingly, production of HA-DHFR-TGA was almost completely ablated in the presence of purified effector and NAD^+^ (**Figure 3A**), with more modest reductions of HA-DHFR-TAA and -TAG. In addition, we detected strong ADP-ribosylation of RF1 and RF2 in reactions where purified effector and NAD^+^ were added. These findings support the notion that effector-Eco9 ADP-ribosylates both RF1 and RF2, but prefers RF2 as a substrate, to disrupt translation.

**Figure 3.**
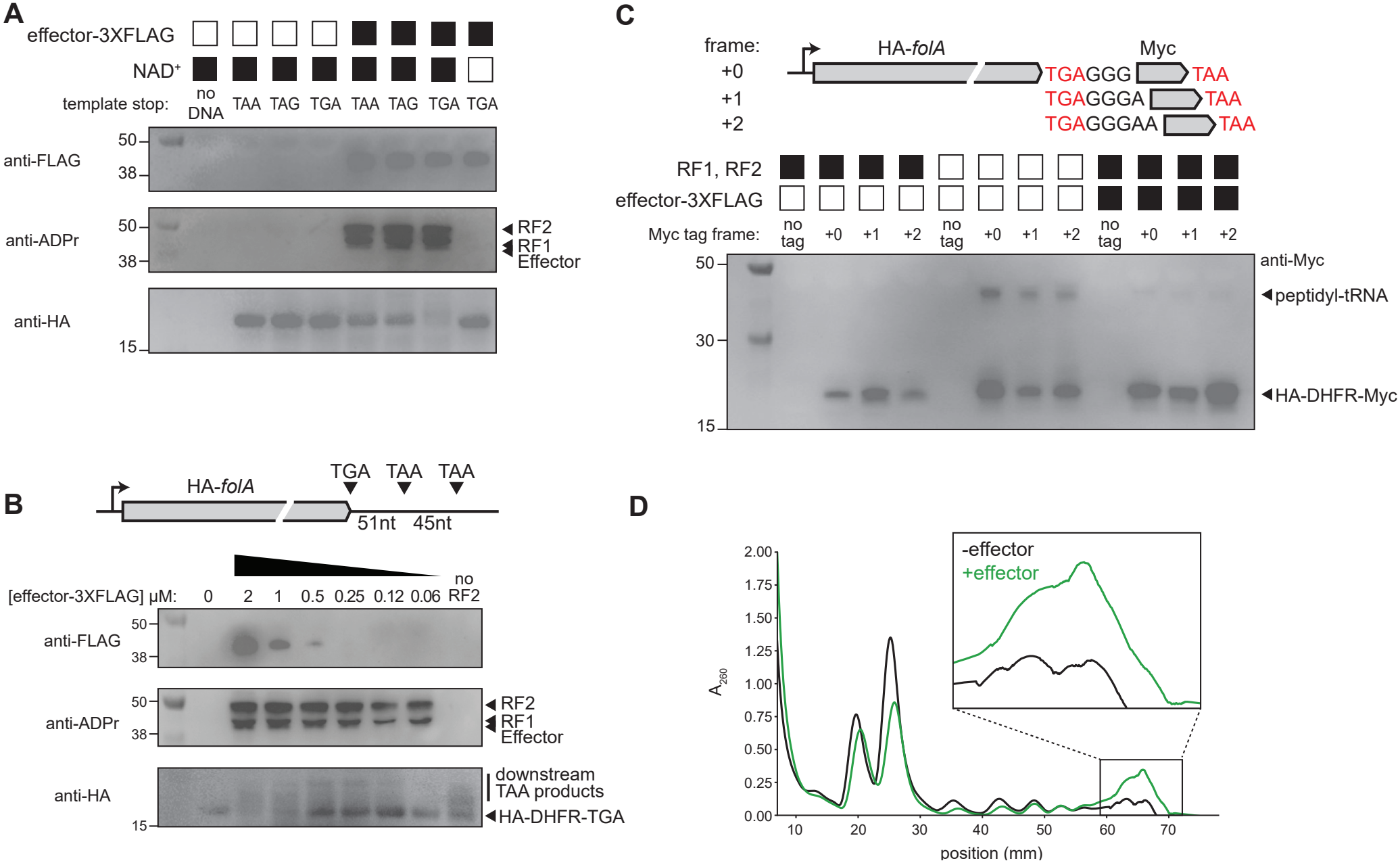
ADP-ribosylated RF1 and RF2 disrupt translation termination. **(A)** IVTT assays using HA-DHFR production from a DNA template as the readout of expression activity in immunoblots. DNA templates were modified to include the indicated stop codons. Purified effector and NAD^+^ were added to the reactions where indicated. **(B)** Top, schematic of HA-*folA* template used in the IVTT reaction. Bottom, IVTT reactions with decreasing concentrations of added effector. NAD^+^ was added to all reactions. The contrast of the HA immunoblot was increased to visualize downstream products. **(C)** Top, schematic of HA-*folA*-Myc templates used in the IVTT reaction. Bottom, IVTT reactions with the indicated Myc reporters and added effector. NAD^+^ was added to all reactions. **(D)** Absorbance spectra from 10-55% (w/v) sucrose gradients loaded with IVTT reactions containing purified effector and HA-DHFR-TAG DNA template.

To further understand the effects of RF2 ADP-ribosylation, we monitored production of HA-DHFR-TGA in the IVTT system with varying concentrations of purified effector or in a control reaction lacking RF2. At effector concentrations where RF1 and RF2 ADP-ribosylation was incomplete, we observed higher molecular weight bands that likely correspond to HA-DHFR products that terminate at TAA stop codons downstream of the usual TGA codon (**Figure 3B**). We also observed higher molecular weight HA-DHFR products in the absence of purified RF2 and purified effector, but these products formed a different banding pattern.

To better understand ribosomal activity at TGA stop codons in the presence of ADP-ribosylated RF2, we modified our original HA-DHFR-TGA template to encode a Myc tag downstream of the TGA stop codon in each possible reading frame. Without added effector, we detected low levels of Myc-tagged reporter (**Figure 3C**). Without RF1 or RF2 in the reaction, these levels increased slightly, and we observed a much higher molecular weight band that likely represents uncleaved peptidyl-tRNA complex as it was depleted following nuclease treatment (**Figure S2G**). With purified effector, we observed minimal signal for the uncleaved peptidyl-tRNA complex and stronger Myc signal, particularly in the +2 frame (**Figure 3C**), suggesting that ribosomes that cannot terminate at TGA using ADP-ribosylated RF2 can frameshift and continue translation until they successfully terminate at a subsequent stop codon. This result also suggests that the activity of modified RF1 and RF2 is distinct from the absence of these release factors completely.

To examine how the distribution of ribosomal particles changed in reactions with purified effector, we loaded an IVTT reaction containing purified effector, NAD^+^, and HA-DHFR-TGA template onto a sucrose gradient after 20 minutes of incubation. Compared to a control reaction lacking effector, we observed a sharp decrease in the monosome fraction as well as increased absorbance at the end of the gradient. Given the increased stop codon readthrough observed (**Figure 3C**), we infer that these are ribosomes accumulating at the ends of transcripts (**Figure 3D**).

Taken together, our IVTT findings support a model where, at low concentrations of effector, RF2 is preferentially ADP-ribosylated relative to RF1, and the modified RF2 cannot faithfully release the ribosome at TGA stop codons. As a result, the ribosome continues until it reaches a TAA stop codon recognized by RF1, switching reading frames in some cases. At high concentrations of effector, both RF1 and RF2 are strongly ADP-ribosylated; without a functional termination pathway ribosomes can complete only a single round of translation such that DHFR does not accumulate to high levels.

### Effector-Eco9 co-opts the bL7/bL12 stalk to associate with the ribosome

To understand how effector-Eco9 targets RF1 and RF2 *in vivo*, we used anti-FLAG beads to immunoprecipitate effector-Eco9(E108A)-FLAG or an untagged version from cell lysates (**Figure S3A**). We used the inactive variant as it may better trap cellular binding partners; we also verified that the C-terminal FLAG tag does not interfere with function (**Figure S3B**). Using tandem MS/MS, we observed substantial enrichment of ribosomal protein bL7/bL12 (bL7 is identical to bL12 but acetylated on its N-terminus) and uL10, which interacts with bL7/bL12 (**Figure 4A**). We reasoned that uL10 was enriched due to its interaction with bL7/bL12, so we focused on bL7/bL12, which comprises the ribosomal stalk and helps recruit translation factors, including EF-G, EF-Tu, IF2, and RF1, RF2, and RF3^22,23^.

**Figure 4.**
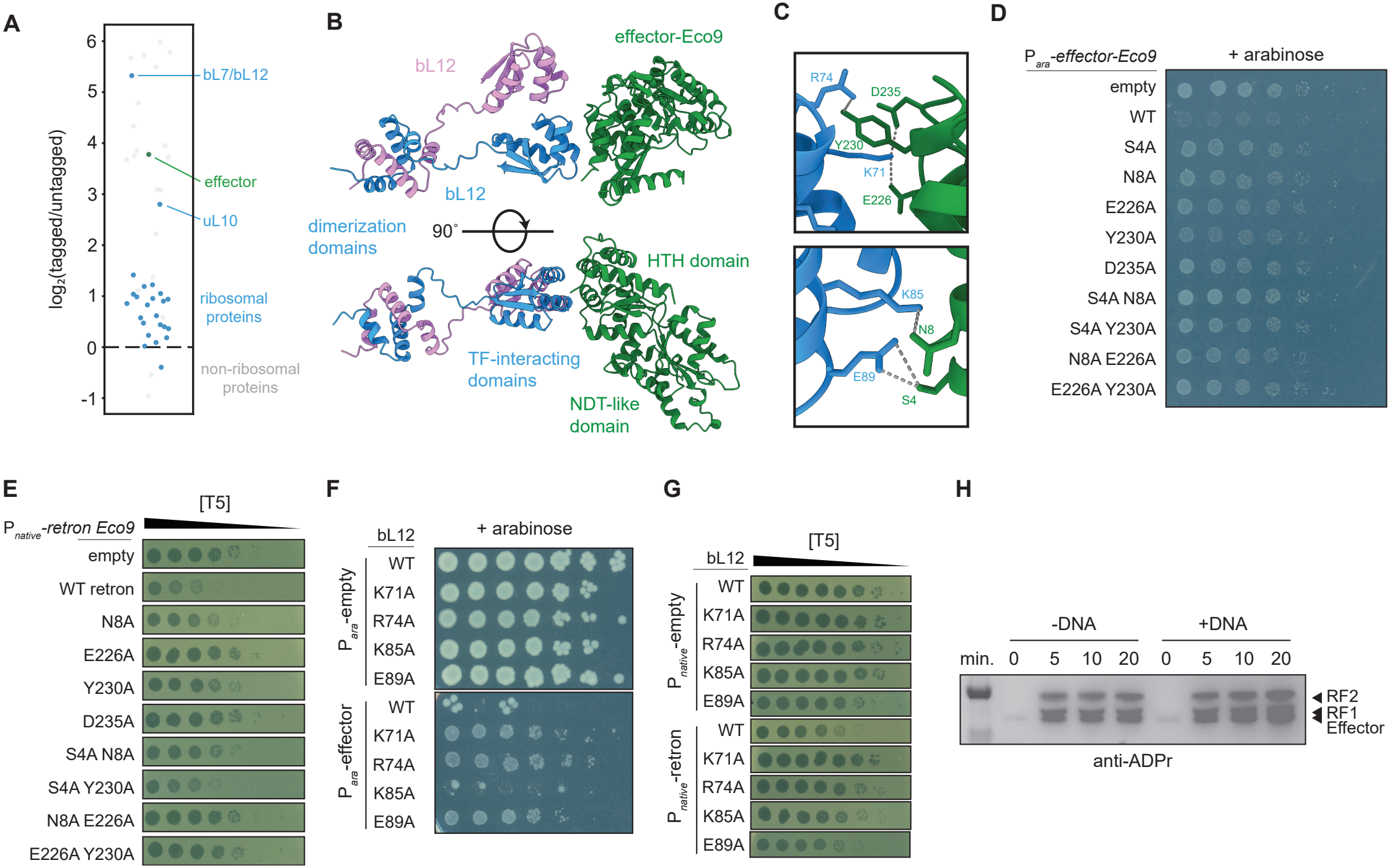
Effector-Eco9 co-opts the bL7/bL12 stalk to associate with the ribosome. **(A)** Mass spectrometry analysis of peptides immunoprecipitated from cells expressing Effector-Eco9(E108A)-FLAG using anti-FLAG beads. The log_2_ ratios of spectral counts for proteins from FLAG-tagged and untagged samples are shown. Proteins with less than 50 spectral counts in the tagged condition were excluded, and a pseudocount was added to facilitate ratio calculations. Ribosomal proteins are labeled in blue, Effector-Eco9 is labeled in green, and all other proteins are labeled in gray. **(B)** AlphaFold3 prediction of a bL7/bL12 dimer (blue and pink) and Effector-Eco9 (green). **(C)** Zoomed view of interacting residues between one bL7/bL12 monomer (blue) and Effector-Eco9 (green) in predicted structure. **(D)** Cell viability assessed by serial dilutions for strains expressing wild-type or mutant versions of Effector-Eco9 from an arabinose-inducible promoter. **(E)** Serial dilutions of T5 phage spotted on lawns of cells containing plasmids expressing wild-type Retron-Eco9 or Retron-Eco9 containing the indicated mutations in Effector-Eco9. **(F)** Cell viability assessed by serial dilutions for strains containing indicated mutations in genomic *rplL* locus (encoding bL7/bL12) and plasmids expressing Effector-Eco9 from an arabinose-inducible promoter or an empty vector. **(G)** Serial dilutions of T5 phage spotted on lawns of cells containing indicated mutations in *rplL* locus and plasmids expressing wild-type Retron-Eco9 or an empty vector. **(H)** IVTT assays containing purified effector, NAD^+^, and HA-DHFR-TAA DNA template where indicated. Timepoints were taken following addition of NAD^+^, and immunoblots were used to assess ADP-ribosylation.

We used AlphaFold3 to predict a complex of effector-Eco9 and a dimer of bL12, which is thought to exist as two dimers independently tethered to the 50S subunit^24,25^ (**Figure 4B**). AlphaFold3 predicted, with high confidence, an interface between the HTH domain of effector-Eco9 and the C-terminal globular domain of one bL12 monomer (**Figure 4B-C, S3C**). The same region in bL12 predicted to bind effector-Eco9 interacts with multiple translation factors^22^, so the residues at this interface are highly conserved.

To test the function of the AF3-predicted interface, we made alanine substitutions in residues of effector-Eco9 predicted to interact with bL12 (**Figure 4D**) and tested whether they reduced effector toxicity. Relative to wild-type effector-Eco9, the variants with S4A, N8A, E226A, Y230A, D235A, or multiple substitutions, each showed enhanced growth in a bacterial spotting assay (**Figure 4D, S3D**). We also introduced the same set of alanine substitutions into effector-Eco9, but in the context of the full retron under the control of its native promoter. We challenged these mutant retron-Eco9 systems with phages T5 and Lust (**Figure 4E, S3E-F**), finding that the single substitutions E226A and D235A completely abolished defense against T5 and Lust, suggesting that the salt bridges formed between E226 and D235 in effector-Eco9 and K71 in bL12 are critical for the interaction. Other alanine substitutions (N8A and Y230A) had more moderate effects on defense, suggesting that these residues play secondary roles in mediating interactions in the protein complex.

To further probe the interaction between effector-Eco9 and bL12, we mutated the chromosomally-encoded bL12 to introduce alanine substitutions in residues predicted to interact with the effector. We then transformed these strains with a plasmid encoding the wild-type effector under the control of an arabinose-inducible promoter. Upon inducing the effector, the K71A, R74A, and E89A variants of bL12 each led to substantially dampened toxicity, relative to wild-type bL12 (**Figure 4F, S3G**). We also transformed the same mutant strains with a plasmid encoding retron-Eco9 under its native promoter and again challenged these strains with the phages T5 and Lust (**Figure 4G, S3H-I**). An alanine substitution at K71 entirely abolished Retron-Eco9 defense against T5, while alanine substitutions at other residues dampened defense against T5 to a moderate degree, except for E89. Taken together, our mutational analyses demonstrate that Eco9 association with the ribosome stalk is necessary for defense.

Considering interactions between the effector and bL12, we reasoned that active translation would potentiate ADP-ribosylation of RF1 and RF2 by recruiting release factors to the ribosome where the effector lies in wait. To test this, we added purified effector to an IVTT reaction with or without DNA template to control translation initiation. In the presence of DNA template, we observed faster ADP-ribosylation of RF1 and RF2 in the IVTT reaction, compared to the reaction lacking DNA (**Figure 4H**). This suggests that effector-bL12 interactions promote ADP-ribosylation of RF1 and RF2.

### Effector-Eco9 increases ribosome occupancy at start and stop codons *in vivo*

To characterize how effector-Eco9 impacts translation *in vivo*, we first examined polysome profiles using sucrose gradients to separate various ribosome particles (**Figure 5A**). Notably, gradients from cells producing effector-Eco9 for 10 or 30 minutes had less distinct polysome peaks relative to an untreated control and a substantial increase in absorbance at the very end of the gradient, indicative of high molecular weight ribosomal species and consistent with IVTT gradients containing purified effector (**Figure 3D**). A similar pattern has been seen in cells with disrupted termination as ribosomes accumulate at the 3’ termini of transcripts^26^.

**Figure 5.**
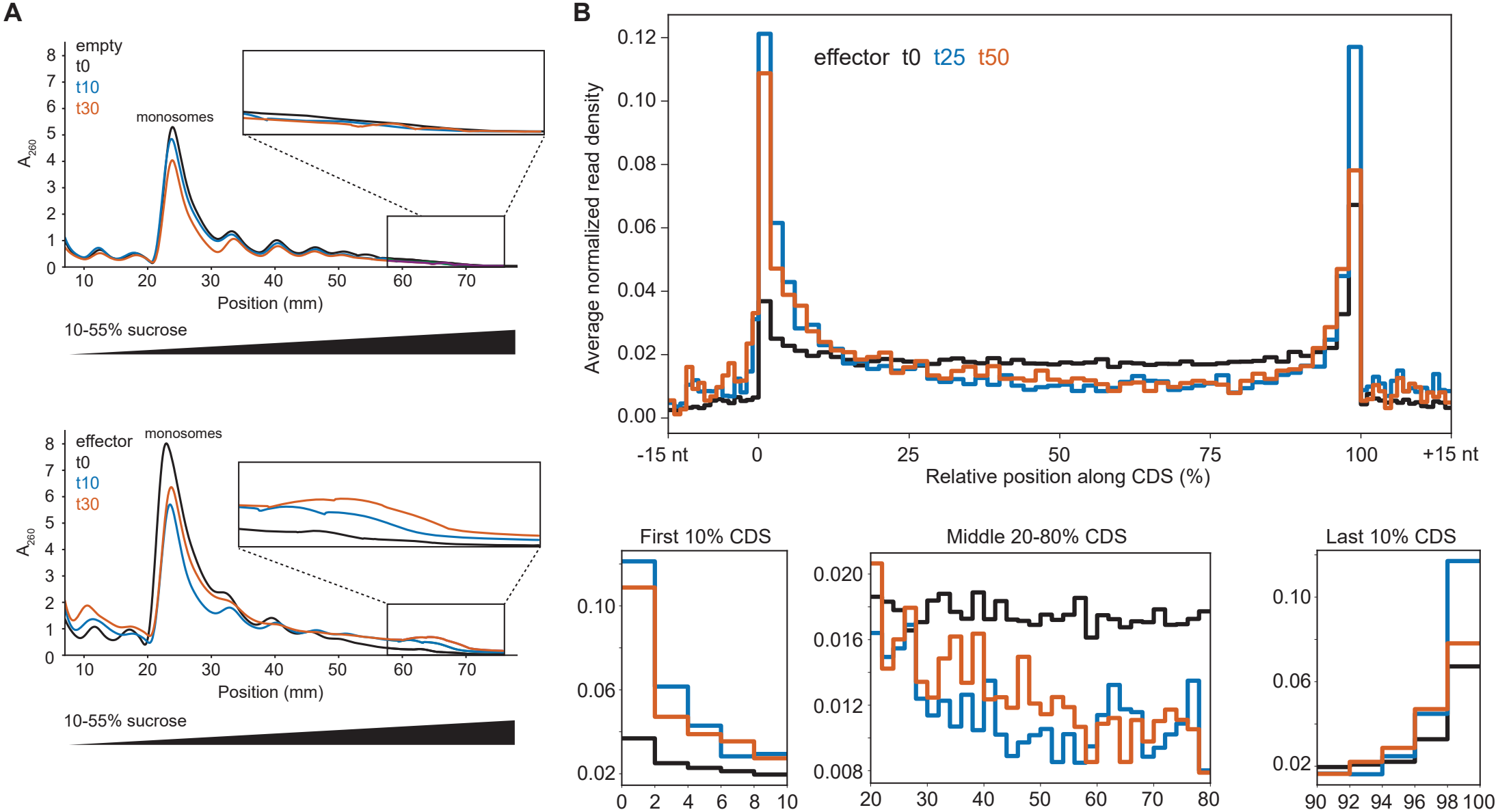
Effector-Eco9 affects ribosome occupancy at start and stop codons *in vivo*. **(A)** 10-55% (w/v) sucrose gradients from lysates harvested from cells expressing effector-Eco9 from an arabinose-inducible promoter at the times indicated post-addition of arabinose. Representative of two biological replicates. **(B)** Average ribosome density across *E. coli* CDSs for cells expressing effector-Eco9 from an arabinose-inducible promoter and harvested at the times indicated post-arabinose addition. Right, zoomed view of first 10 percent, middle 20 to 80 percent, and last 10 percent of CDSs.

To further probe how effector-Eco9 inhibits translation, we used ribosome profiling to map and quantify ribosome-protected transcripts after producing effector-Eco9 for 0, 25, or 50 minutes. Visual inspection of ribosome occupancy across individual transcripts suggested that ribosome density across coding regions generally decreased at 25 and 50 min post-induction, with peaks remaining at start and stop codons. To quantify these trends, we performed a metagene analysis wherein we calculated the normalized fraction of reads across 50 equally sized bins for each coding sequence and then averaged those read profiles for all coding sequences. At 25 minutes post-induction, we observed a roughly 3-fold increase in reads around start codons and an approximate 2-fold increase in reads around stop codons (**Figure 5B**). These increases were accompanied by a substantial decrease in occupancy throughout the rest of the coding region with occupancy more reduced toward the 3’ end of transcripts. Similar patterns were seen at 50 minutes post-induction, though without the increase in occupancy around stop codons.

Next, we performed cryogenic electron microscopy (cryoEM) on cells producing effector-Eco9 or an empty vector. We used cryoPRISM^27^ (see Methods), which images ribosomes directly from cell lysates, *i*.*e*., without purifications. We obtained cryoEM density maps of polysomes, monosomes, and dissociated large and small subunits; cryoDRGN and related tools^28–31^ were then used to categorize sub-populations within each ribosomal class (**Figure S4-S8**). We observed an increase in 70S ribosomes relative to dissociated ribosomal subunits in the effector-producing cells (**Figure 6A**, left). Notably, we could resolve mRNA and P-site tRNA on all 70S ribosomes in effector-producing cells, whereas without the effector 47% of 70S ribosomes occupied an idle, translationally inactive state, *i*.*e*., without mRNA or P-tRNA.

**Figure 6.**
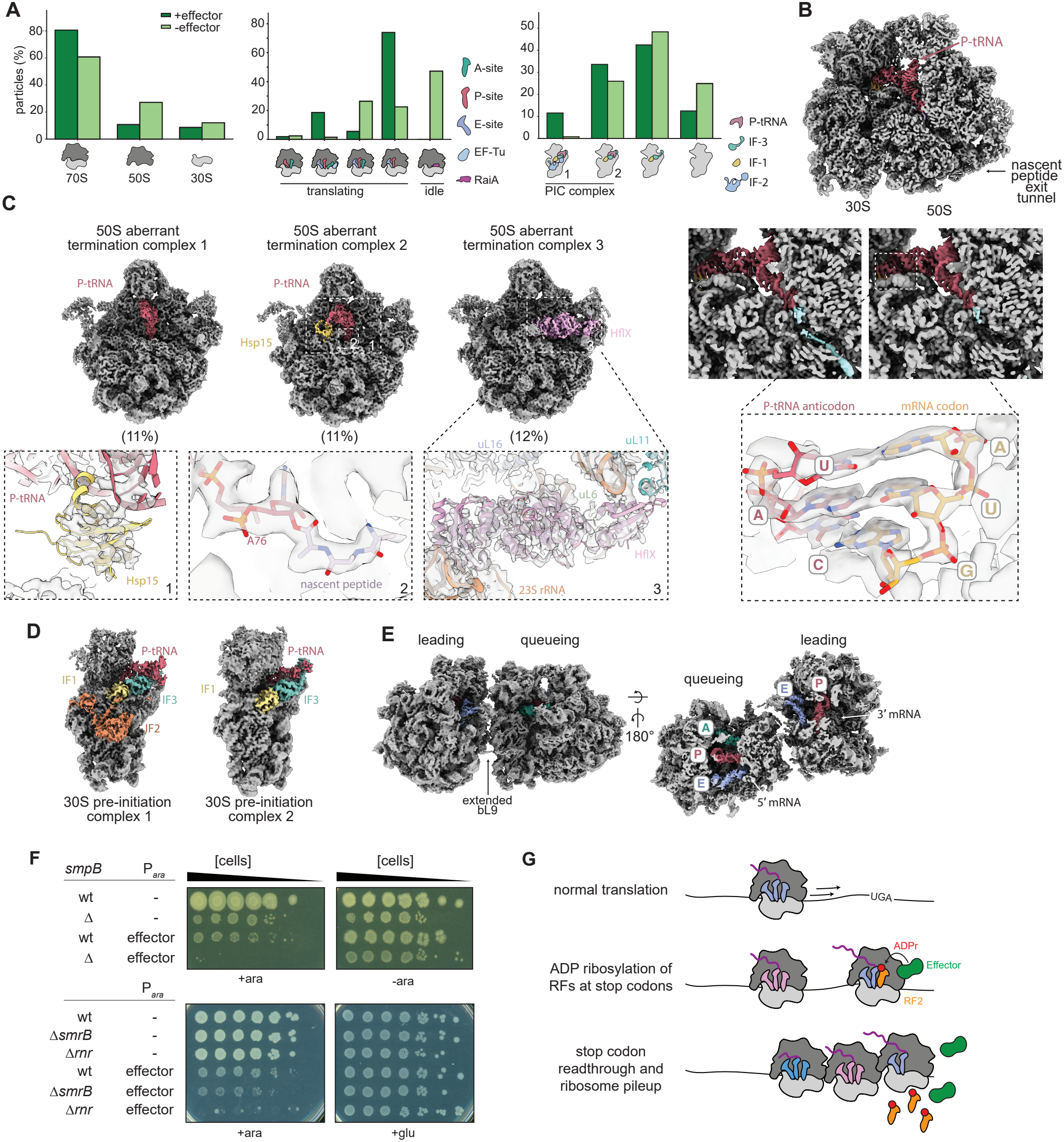
Effector-Eco9 disrupts translation termination and triggers ribosome rescue pathways. **(A)** Distribution of 70S, 50S, and 30S ribosomal particles identified by cryoEM from cells expressing effector-Eco9 or containing an empty vector (left), 70S sub-states (middle), and 30S sub-states (right). **(B)** Density map of a 70S ribosome from cells expressing effector-Eco9, containing P-site tRNA (red) and an empty A-site. The map is clipped to show the nascent peptide exit tunnel (NPET). 70S particles were further classified based on the presence (top left inset) or absence (top right inset) of density (cyan) within the NPET. For the latter class, an atomic model (pdb 7st6)^65^ for an AUG codon-anticodon pairing was fit into the density at the decoding center (bottom inset). **(C)** Density maps for three 50S subunit complexes observed only in cells producing effector-Eco9. Aberrant termination complexes 1 (left) and 2 (middle) contain P-site peptidyl-tRNA (red) and density extending from the terminal nucleotide of the CCA acceptor end, into which a short nascent peptide was modelled, as shown for complex 2 (inset 2). Complex 2 contains additional density consistent with Hsp15, into which an atomic model (pdb 8ap4)^66^ was docked (inset 1). Complex 3 contains density for the stress-dependent ribosome splitting factor HflX (pdb 5ady)^67^ (inset 3). All models were refined using ISOLDE^68^. The % of 50S subunits assigned to each complex is indicated (*overleaf*). **(D)** Density maps for two 30S pre-initiation complexes (PICs) enriched in cells producing effector-Eco9. PIC complex 1 contains IF1, IF2, and IF3 and P-site tRNA (pdb 9h9h; 5lmq)^69–70^ and accounts for 12% of 30S subunits. PIC complex 2 contains IF1, IF3, and P-site tRNA (pdb 5lmq) and accounts for 34% of 30S subunits. **(E)** Density maps for disomes comprised of a leading and a queueing ribosome from cells producing effector-Eco9, shown in a front (left) and top-down clipped (right) view. The leading ribosome exhibits an extended bL9 conformation that contacts the small subunit of the queueing ribosome. The clipped density map (right) shows A, P, and E-site tRNAs in the queueing ribosome, whereas the A-site of the leading ribosome lacks tRNA. **(F)** Cell viability assessed by serial dilutions for strains containing indicated genomic deletions and effector-Eco9 under the control of an arabinose-inducible promoter or an empty vector. Top panel is LB medium, bottom is M9L medium. Both panels in this figure are representative of two replicates. **(G)** Schematic of Effector-Eco9 targeting and its effects on ribosomes and translation termination and fidelity.

Additionally, in effector-producing cells, we observed a ~4-fold decrease in particles with tRNA present in the A, P, and E-sites, with a concomitant increase in particles bearing only P- and E-site tRNAs (**Figure 6A**, middle). We separated these particles into two similarly sized classes distinguished by the presence or absence of density inside the nascent peptide exit tunnel (NPET). Ribosomes with density inside the tunnel allowed us to build a nascent chain spanning most of the tunnel’s length (**Figure 6B**, top left inset), and are consistent with particles stalled during translation at a step preceding termination. The other class of particles lacked density in the NPET (**Figure 6B**, top right inset), and exhibited codon-anticodon pairing consistent with an AUG start codon, suggesting they are stalled at initiation, though we could not unambiguously assign the bases (**Figure 6B**, bottom inset). Collectively, these analyses were consistent with our ribosome profiling data indicating that the effector leads to an accumulation of ribosomes at start and stop codons.

### Effector-Eco9 disrupts translation termination *in vivo*

In the IVTT reactions, effector-Eco9 disrupted translation termination (**Figure 2A, 3A-D**), so we examined our cryoEM datasets for indications of aberrant translation. We first inspected dissociated large subunits. Despite the smaller fraction of large subunits in the effector-treated cells (**Figure 6A**, left), three unique ribosomal complexes emerged (**Figure 6C, S6D, S7D-F**), each consistent with aberrant termination. In complexes 1 and 2 we observed a fully assembled 50S subunit bound by P-site peptidyl-tRNA, with complex 2 additionally exhibiting density near a known binding site for Hsp15 (**Figure 6C**, inset 1), which has been implicated in rescuing ribosomes stalled during termination^32–34^. Inspection of the CCA acceptor end of these P-site tRNAs revealed clear density extending from the terminal nucleotide, confirming that this complex arises without canonical termination and hydrolysis of the peptidyl-tRNA linkage (**Figure 6C**, inset 2). Complex 3 contained density corresponding to the ribosome splitting factor HflX, a GTPase implicated in rescuing stalled ribosomes (**Figure 6C**, inset 3)^35,36^. The emergence of these three post-splitting complexes suggests a failure to properly terminate translation in cells producing effector-Eco9.

Given that roughly one-third of dissociated 50S subunits were undergoing ribosomal rescue and recycling following effector production, we reasoned that these particles would be unavailable for binding to initiation-competent small subunits, leading to an accumulation of such states. Consistent with this hypothesis, we detected an enrichment of two pre-initiation complexes compared to our control – one containing IF1, IF2, IF3, and P-site tRNA and another bearing IF1, IF3, and P-site tRNA (**Figure 6A, 6D, S6E, S8A-B**).

Finally, because our polysome profiles (**Figure 3D, 5A**) revealed very high molecular weight polysomes following effector-Eco9 production, a phenotype seen with other termination defects^37–39^, we were curious whether these higher-order ribosome assemblies bore distinctive structural features. Disome complexes, including those awaiting termination, can assemble when the leading ribosome stalls, causing the queuing ribosome to collide. Such complexes can be distinguished from actively translating particles based on the absence of A-site tRNA in the leading particle. Among our 70S ribosome classes, we observed a subset of ribosomes that, after particle extraction with larger box size (see Methods), allowed us to resolve disome pairs bearing a characteristic inter-ribosome contact mediated by bL9 (**Figure 6E, S6C, S8C-D**). For cells producing effector-Eco9, the leading ribosome had E- and P-site tRNAs but typically lacked A-site tRNA, whereas the queuing ribosome usually had all three tRNAs. In contrast, untreated cells had occupied A-sites on both leading and queueing ribosomes. The stalled/collided complexes in cells producing effector-Eco9 may represent ribosomes that reached the 3’ ends of mRNAs without having terminated at the encoded stop codons; the 3’ end of the mRNA would sit in the P-site, leaving the A-site free of both mRNA and tRNA, as observed. These analyses further support a model in which effector-Eco9 disrupts translation termination.

### ADP-ribosylation of RF1 and RF2 triggers ribosome rescue pathways

To complement these studies, we performed cryoEM on IVTT reactions containing purified effector, NAD^+^, and HA-DHFR-TGA template, using a similar analysis pipeline. Like our *in vivo* studies, in effector-treated samples, we commonly observed ribosomal states characteristic of translation termination failure (**Figure S9-S11**), including an enrichment of disomes with the leading ribosome lacking an A-site tRNA (**Figure S12**). Notably, we did not observe states indicative of active ribosome rescue, as IVTT reactions lack the ribosome rescue pathways found *in vivo* that work to resolve stalled ribosomes and degrade the associated mRNA and aberrant protein. This cellular degradation of mRNA would explain the 3’-5’ bias in ribosome occupancy seen by ribosome profiling (**Figure 5B**), as well as the ribosomes stalled at initiation codons, which would protect the 5’ end of mRNAs from degradation.

In sum, we hypothesized that the difference in ribosome states seen *in vitro* and *in vivo* reflects the role that ribosome rescue plays in cells following effector-Eco9 production. This model predicts that effector toxicity would be exacerbated by the absence of the primary ribosome rescue pathway consisting of tmRNA, SmpB, and RNaseR^40^. Further, because our cryoEM and sucrose gradient studies indicated that stalled/collided ribosomes may accumulate at the 3’ termini of mRNAs, we also sought to assess effector toxicity in cells lacking SmrB, the endoribonuclease that helps rescue collided ribosomes^41^. Strikingly, we found that producing effector-Eco9 at low levels in *ΔsmpB* and *Δrnr* cells produced almost no viable colonies, compared to wild-type cells expressing effector-Eco9 at the same levels (**Figure 6F**). In *ΔsmrB* cells there was a modest increase in effector toxicity. These results help to explain the difference in ribosome fates *in vitro* and *in vivo* following effector-Eco9 induction.

For *in vitro* reactions, ribosomes accumulate at the ends of mRNAs, whereas *in vivo*, ribosome rescue pathways likely act to try and salvage such stalled ribosomes, producing the transient accumulation of ribosomes at stop codons seen by ribosome profiling as well as the aberrant ribosome particles seen by cryoEM that stem from perturbed termination and ribosome rescue.

## DISCUSSION

Across the diverse collection of characterized anti-phage defense systems, translation stands as a prominent molecular target. Anti-phage defense systems like RAZR^42^, PARIS^43^, and numerous retron systems (Eco2^44^, Eco7^45^, Kva2^46,47^, Sen3^48^, among others) provide robust defense by targeting translation-associated nucleic acids. Here, we demonstrate that Retron-Eco9 also ablates translation, but does so by ADP-ribosylating the essential translation termination factors RF1 and RF2, which have not previously been revealed as the targets of a defense system. To date, the only other defense system that directly targets a translation factor is the Lit protease, which cleaves EF-Tu to halt T4 infection^12^. However, there may be other examples. For instance, Doc toxins from Doc-PhD toxin-antitoxin systems phosphorylate EF-Tu^49^; while Doc-PhD systems have not been reported as anti-phage, the stand-alone Doc-like protein SoFic can provide anti-phage defense^50^, but its molecular target remains unknown.

Effector-Eco9 appears to impact translation through a cascading effect (**Figure 6G**); ADP-ribosylated RF1 and RF2 disrupt termination, but can also increase stop codon readthrough and frameshifting. As ribosomes accumulate and collide, likely at the ends of mRNAs, ribosome rescue pathways try to salvage ribosomes, leading to the aberrant 50S particles documented by cryoEM. Accumulating such defective 50S particles may limit the formation of new initiation complexes, leading to the observed 30S particles bound to IF2, an intermediate not seen in unperturbed cells. We speculate that other translation-targeting defense systems may similarly trigger complex cellular responses.

Effector-Eco9 joins several other ADP-ribosyltransferases in anti-phage defense, including CmdT^9^, DarT^51^, and Effector-Eco1^14^. ADP-ribosyltransferases are also commonly used as effectors in type VI secretion systems, including RhsP2^52^, which targets RNA, and Tre^Tu^, which ADP-ribosylates EF-Tu in other bacterial cells^53^. Important bacterial pathogens also commonly secrete ADP-ribosyltransferase toxins to inhibit host eukaryotic cells, including the cholera toxin from *Vibrio cholerae*, which ADP-ribosylates G-proteins^54^, and diphtheria toxin from *Corynebacterium diphtheriae*, which modifies eEF2 to block translation^55^. Taken together, these various toxins and defense systems highlight a major role for ADP-ribosylation as a molecular mechanism that bacteria use to manipulate and impede cellular processes.

Histidine ADP-ribosylation of proteins remains poorly studied relative to other amino acids, though there are some examples. Diptheria toxin ADP-ribosylates the imidizole ring in dipthamide, a modified histidine. In plants and bacteria, some TIR domains ADP-ribosylate histidine that is not incorportated into proteins, using His-ADPR as an immunity signaling molecule^56^. There are also proteomic studies indicating that histidine ADP-ribosylation may be common in mammals^57^, but the enzymes responsible for it are not clear. Our work suggests that some NDT-like enzymes in addition to traditional ADP-ribosyltransferases may be important ‘writers’ of this modification.

## METHODS

### Strains and growth conditions

Bacterial strains and phages used in this study are listed in **Supplementary File 1**. *Escherichia coli* strains were routinely cultured in Luria broth (LB) medium at 37 °C for maintenance and cloning purposes. Phage amplification was carried out by infecting *E. coli* MG1655 cultures at an OD_600_ of 0.1-0.2 using a multiplicity of infection (MOI) of ~0.1. Following lysis, cultures were centrifuged to remove bacterial debris, and the supernatant was passed through a 0.2 µM filter. All phage infection experiments in liquid culture and phage spotting assays were performed in LB medium at 25 °C. For incorporation assays, bacterial cultures were grown in M9 minimal media (6.4 g/L Na_2_HPO_4_-7H_2_O, 1.5 g/L KH_2_PO_4_, 0.25 g/L NaCl, 0.5 g/L NH_4_Cl) supplemented with 0.1% casamino acids, 0.4% glycerol, 0.4% glucose, 2 mM MgSO_4_, and 0.1 mM CaCl_2_ at 37 °C. All other experiments were performed in LB medium at 37 °C. Antibiotics were added as needed for plasmid selection and maintenance at the following final concentrations (liquid; plates): carbenicillin (50 µg/mL; 100 µg/mL), chloramphenicol (20 µg/mL; 30 µg/ mL), kanamycin (30 µg/mL; 50 µg/mL), and spectinomycin (50 µg/mL; 50 µg/mL).

Yeast strains used in this study are listed in **Supplementary File 1**. *S. cerevisiae* strains were routinely cultured in YPD medium at 30 °C for maintenance and cloning purposes. Antibiotics were added as needed for plasmid selection and maintenance at the following final concentrations (plates): G418 (200 mg/L), nourseothricin (100 mg/L).

### Plasmid construction

All plasmids, primers, and gBlocks are detailed in **Supplementary File 1**.

#### pBAD33::Effector-Eco9 constructs

Effector-Eco9 is present in the genome assembly of *E. coli* NILS-16 (NCBI Assembly accession: GCA_014461615.1), on contig DABRBC010000011, at positions 2488-3474 on the reverse strand. The effector was amplified from pJB145 (pSC101::Retron-Eco9) with primers CB139 and CB140 and inserted into pBAD33 linearized with primers CB45 and CB46 using Gibson assembly. pBAD33::Effector-Eco9(E108A) was constructed via around-the-horn PCR using CB154 and CB155 primers and pCB48 (pBAD33::Effector-Eco9) as template. Additional pBAD33::Effector-Eco9 variants with mutations at the bL7/ bL12 interface (S4A, N8A, E226A, Y230A, D235A, and double mutants) were constructed using Gibson assembly. Insert fragments were amplified from pCB48 with primers CB140 and CB620 (S4A), CB140 and CB388 (N8A), CB140 and CB392 (Y230A), CB140 and CB390 (E226A), CB46 and CB622 (D235A). Backbone fragments were amplified from pCB48 with primers CB45 and CB621 (S4A), CB45 and CB389 (N8A), CB45 and CB393 (Y230A), CB45 and CB391 (E226A), and CB139 and CB623 (D235A). For double mutant constructs, insert and backbone fragments were amplified from pCB155 (S4A N8A, N8A E226A), pCB156 (S4A Y230A), or pCB157 (E226A Y230A). Insert fragments were amplified with primers CB140 and CB620 (S4A N8A, S4A Y230A), CB140 and CB390 (N8A E226A), and CB140 and CB392 (E226A Y230A). Backbone fragments were amplified with primers CB45 and CB621 (S4A N8A, S4A Y230A), CB45 and CB391 (N8A E226A), and CB45 and CB393 (E226A Y230A).

pBAD33::empty and pBAD::Effector-Eco9 plasmids containing carb^R^ cassettes were constructed by Gibson assembly. Primers CB692 and CB693 were used to amplify the carb^R^ cassette from pCB1 (pBR322::empty), and primers CB690 and CB691 were used to amplify the pBAD33 backbone from pCB120 (pBAD33::empty) or pCB48.

pBAD33-pMB1 origin plasmids (empty, Effector-Eco9, Effector-Eco9(E108A), and Effector-Eco9(E108A)-FLAG) were constructed by Gibson assembly. Primers CB325 and CB326 were used to amplify the pMB1 origin from pCB1, and primers CB324 and CB336 were used to amplify the pBAD33 backbone from pCB120 (empty), pCB48 (Effector-Eco9), pCB55 (Effector-Eco9(E108A)), or pCB142 (Effector-Eco9(E108A)-FLAG). pBAD33-pMB1 origin::Effector-Eco9-FLAG was constructed by around-the-horn PCR using primers CB327 and CB328 and pCB136 as template.

#### pSC101::Retron-Eco9 constructs

All plasmids were constructed by Gibson assembly. All insert fragments were amplified using primers CB457 and CB458, except for the Y230A insert which was amplified using primers CB393 and CB457. Insert fragments were amplified using the following templates: pCB155 (N8A), pJB145 (Y230A), pCB157 (E226A), pCB253 (D235A), pCB255 (S4A N8A), pCB256 (S4A Y230A), pCB159 (N8A E226A), and pCB160 (E226A Y230A). All backbone fragments were amplified using primers CB203 and CB459, except for the Y230A backbone which was amplified using primers CB203 and CB392. All backbone fragments were amplified from pJB145.

#### pT7::*folA* constructs

pT7::*folA*-GSSG-FLAG was constructed by around-the-horn PCR using primers CB247 and CB248 and the pT7::*folA* template from the PURExpress reconstituted protein synthesis system (NEB). To construct pT7::*folA* plasmids with different stop codons, the TAA stop codon in pT7::*folA* was first changed to TAG or TGA by around-the-horn PCR using primers CB883 and CB884 (TAG) and CB883 and CB885 (TGA) and pT7::*folA* as template. Next, primers CB892 and CB877 were used to add a N-terminal HA-tag using around-the-horn PCR and pT7::*folA*-TAA, -TAG, or -TGA as template. To construct pT7::*folA*-TGA plasmids with downstream Myc tags in different reading frames, around-the-horn PCR was used with pCB362 (pT7::*folA*-TGA) as template. Primers were used to insert the Myc tag in one of three reading frames: CB938 and CB939 (+0 frame), CB938 and CB940 (+1 frame), CB938 and CB941 (+2 frame).

#### Plasmids for *rplL* genomic mutations

pVan::*rplL* was constructed by Gibson assembly; primers CB384 and CB385 were used to amplify the insert fragment from the MG1655 genome, and primers CB308 and CB309 were used to amplify the pVan backbone from pCB121 (pVan::empty). Mutations in *rplL* were introduced using around-the-horn PCR and pCB162 (pVan::*rplL*) as template. Primers CB487 and CB489 (K71A), CB490 and CB492 (R74A), CB493 and CB495 (K85A), and CB626 and CB627 (E89A) were used to introduce the indicated mutations. pKDsgRNA plasmids were constructed using around-the-horn PCR from pCB203 (pKDsgRNA::p15a) template DNA and primers CB535 and CB538 (K71), CB536 and CB538 (R74), CB537 and CB538 (K85), and CB628 and CB538 (E89).

#### *prfA* and *prfB* constructs

pVan::*prfA* (pCB244) and pVan::*prfB* (pCB245) were constructed by Gibson assembly. Insert fragments were amplified from MG1655 using primers CB596 and CB597 (*prfA*) and CB598 and CB599 (*prfB*), and backbone fragments were amplified from pCB121 using primers CB308 and CB309. C-terminal GSSG-3XFLAG fragments were added to each plasmid by around-the-horn PCR using pCB244 or pCB245 as template and primers CB860 and CB927 (*prfA*-GSSG-3XFLAG) and CB860 and CB861 (*prfB*-GSSG-3XFLAG). The H253A mutant was introduced into pCB345 (pVan::*prfB*-GSSG-3XFLAG) by around-the-horn PCR using primers CB872 and CB873.

#### Additional constructs

ProD::*tufA*-GSSG-3XFLAG was constructed by Gibson assembly and around-the-horn PCR. *TufA* was first amplified from MG1655 using primers CB837 and CB838 and inserted into the ProD backbone amplified by primers CB823 and CB824. The C-terminal GSSG-3XFLAG fragment was then inserted by around-the-horn PCR using primers CB823 and CB824.

pBR322::Retron-Eco9 was constructed by Gibson assembly; the insert fragment was amplified from pJB145 using primers CB156 and CB157 and the backbone fragment was amplified from pCB1 (pBR322::empty) by primers CB1 and CB2. The E108A mutation was introduced into the effector in pCB76 (pBR322::Retron-Eco9) using primers CB154 and CB155 in around-the-horn PCR.

pBAD33::*dam* was constructed by Gibson assembly; the insert fragment was amplified from MG1655 using primers CB133 and CB134 and the backbone fragment was amplified from pCB120 using primers CB45 and CB46.

pBR322::Retron-Eco1 was constructed by Gibson assembly; the backbone fragment was amplified from pCB1 using primer CB1 and CB2 and the insert was ordered as a gBlock (gCB1) from IDT.

### Strain construction

Plasmids described above were introduced into *E. coli* MG1655 or BW27783 by TSS transformation or electroporation.

Genomic *rplL* (encoding bL7/bL12) mutations were generated using the no-SCAR method^58^ as described. Briefly, sequences for RNA guides to target wildtype *rplL* were designed manually, following the no-SCAR guidelines, and selected for targeting wild-type *rplL* but nowhere else in the *E. coli* genome. The guides were inserted into the pKDsg plasmid, which also contains lambda red recombineering machinery, and tested for their ability to target the *E. coli* BW27783 genome. Efficient guides were selected, and the corresponding pKDsg plasmid was co-transformed into *E. coli* BW27783 with the pCas9 plasmid. dsDNA containing the desired mutation in *rplL* was generated by PCR using plasmids containing the corresponding mutant version of *rplL* as template, treated with DpnI at 37 °C, and purified using the Zymo Clean & Concentrate Kit. Strains containing both pKDsg and pCas9 plasmids were made electrocompetent and transformed with dsDNA containing the desired mutation in *rplL*. Cells were then plated on media containing 1X aTc to induce expression of Cas9 and the guide RNA present in the pKDsg plasmid. Colonies were screened by PCR to identify hits with the desired mutation in *rplL*, and both pCas9 and pKDsg plasmids were cured from the final mutant strains.

*SmpB, smrB, rnr, tufA*, and *tufB* deletion strains were constructed by P1 phage transduction from the Keio collection into MG1655. Donor P1 phage was created by combining 100 µL P1 phage with 300 µL turbid donor culture and 5 mM CaCl_2_. After incubation at 37 °C for 20 minutes, the mixture was expanded into 5 mL LB + 5 mM CaCl_2_ and incubated with shaking at 37 °C for 4-6 hours. Cleared cultures were then spun down to remove debris and stored with chloroform until use. 1 mL recipient culture (MG1655) supplemented with 5 mM CaCl_2_ was then pelleted and resuspended in 100 µL volume. 1-50 µL donor P1 phage were added to the recipient culture and incubated for 30 minutes at 37 °C. Cells were then spun down, resuspended in 1 mL LB + 0.2 mM sodium citrate, and incubated for 1 hour at 37 °C shaking. Cells were then plated on LB + kanamycin (all Keio deletions are kan^R^) plates spread with 1M sodium citrate. Single colonies were restreaked to remove remaining phage, and deletions were confirmed by PCR and sequencing.

Δ*tufA* Δ*tufB* ProD::*tufA*-GSSG-3XFLAG was constructed in three parts. First, Δ*tufA* was introduced into MG1655 by P1 transduction. Next, ProD::*tufA*-GSSG-3XFLAG was transformed into the Δ*tufA* strain. Finally, Δ*tufB* was removed by P1 transduction.

GAL1,10-Effector-Eco9-TEV-3XFLAG and GAL1,10-Effector-Eco9(E108A)-TEV-3XFLAG were individually integrated into chromosome X of the *S. cerevisiae* genome at nucleotide position 223616, following the EasyClone-MarkerFree 2.0 system^59,60^. The linear DNA used for integration was first generated by a series of SOE PCRs. gBlocks containing a codon-optimized version of GAL1,10-Effector-Eco9-TEV-3XFLAG-CYC1 terminator (gCB2 and gCB3) were ordered from IDT and contained 40 bps of overlapping sequence. The native GTG start in Effector-Eco9 was also replaced with ATG in gCB2. These fragments were first combined through PCR using primers CB629 and CB630. This PCR product was further amplified using primers CB649 and CB651 to generate fragment B, containing overhangs to additional fragments on either end. Fragment A, containing upstream homology to chromosome X, was amplified from pCfB2189 using primers CB652 and CB653. Fragment C, containing the LEU2 marker and downstream homology to chromosome X, was amplified from pCfB2189 using primer CB650 and CB654. For cloning the E108A mutant, fragment B was amplified with primers CB651 and CB695 to generate fragment B.1 and primers CB649 and CB694 to generate fragment B.2.

Competent cells (containing the pCfB2312 Cas9 expression vector) were prepared following the EasyClone-MarkerFree protocol, and fragments A, B, and C (or A, B.1, B.2, and C for the E108A mutant) were transformed into cells alongside 0.5 µg pCfB3041 (gRNA helper plasmid), 26% (w/v) PEG 3350, 78 mM lithium acetate, and 0.2 mg/mL salmon sperm DNA. Following recovery, cells were plated on YPD agar containing 200 mg/L G418 and 100 mg/L nourseothricin and grown at 30 °C. Colonies were restreaked onto SD-Leu plates and again grown at 30 °C. Finally, individual colonies were replica plated onto YPD + G418 and YPD + nourseothricin to remove Cas9 expression and gRNA helper plasmids. Colonies that did not grow on G418 and nourseothricin were then sequence verified. The wild-type Effector-Eco9 strain contains a synonymous mutation at position P200.

### Phage spotting assays and EOP measurements

To determine phage titers, serial dilutions of phage stocks were mixed with *E. coli* MG1655 and molten LB + 0.5% agar and spread on LB + 1.2% agar plates and incubated at 37 °C overnight. For phage spotting assays, 40 µL of a bacterial strain of interest was combined with 4 mL molten LB + 0.5% agar and overlaid on an LB + 1.2% agar + antibiotic plate. Phage stocks were serially diluted in 1X FM buffer (20 mM Tris-HCl pH 7.4, 100 mM NaCl, 10 mM MgSO_4_), and 2 µL of each dilution was spotted on the bacterial lawn. Plates were incubated at 25 °C overnight before imaging. The efficiency of plaquing (EOP) was determined by comparing plaque formation on the experimental strain relative to the control strain. Experiments were replicated independently three times and representative images are shown.

### Growth curves following phage infection in liquid culture

Single colonies of *E. coli* MG1655 containing either pTU175 (pSC101::empty) or pJB145 (pSC101::Retron-Eco9) were cultured overnight in LB + antibiotics. Overnight cultures were diluted 1:100 in fresh LB + antibiotics and grown at 37 °C until OD_600_ = ~0.2 was reached. Cultures were then further diluted to an OD_600_ of 0.1, and 1 mL of the resulting suspension was transferred into each well of a 24-well plate. T5 or Lust phage were added to each well at the specified MOI, and bacterial growth following infection was measured at 15-minute intervals with orbital shaking at 25 °C using a Biotek plate reader. Data reported are the mean and standard deviation of four technical replicates collected across two biological replicates.

### Bacterial spotting assays

For producing Effector-Eco9 and Effector-Eco9 E108A, single colonies of *E. coli* MG1655 harboring pBAD33::Effector-Eco9 and pBAD33::Effector-Eco9 E108A or the corresponding empty vector were cultured for 6-8 hours at 37 °C in LB + 0.4% glucose + antibiotics. A 300 µL aliquot of each saturated culture was pelleted by centrifugation at 4,000 x *g* for 5 minutes, washed once in 1X phosphate-buffered saline (PBS), and resuspend in 1X PBS to an OD_600_ of 1.0. Cultures were then serially diluted tenfold in 1X PBS, and 3 uL of each dilution was spotted on M9L plates (M9 medium supplemented with 10% LB (v/v)) further supplemented with 0.4% glucose (w/v) or 0.2% arabinose (w/v). Plates were then incubated at 37 °C overnight before imaging.

For producing Effector-Eco9 in the presence of different antibiotics, *E. coli* BW27783 containing pBAD33::Effector-Eco9-carb^R^ or the corresponding empty vector were grown to saturation and processed as above. Cultures were spotted onto M9L plates containing 0.4% glucose (w/v) and 0.6 µg/mL kanamycin, 5 µg/mL retapamulin or 0.25 µg/mL rifampicin, or 2×10^−6^% arabinose (w/v) and the same set of antibiotics. Plates were then incubated at 37 °C overnight before imaging.

To produce Effector-Eco9, *E. coli* BW27783 containing pBAD33::Effector-Eco9 or a mutant form of Effector-Eco9 were grown to saturation and processed as above. Cultures were plated onto 0.4% glucose (w/v) and 2×10^−5^% arabinose (w/v) and incubated o/n at 37 °C.

To produce Effector-Eco9 in bL7/bL12 mutant strains, *E. coli* BW27783 containing mutations in the *rplL* gene and containing pBAD33::Effector-Eco9 or a corresponding empty vector were grown to saturation and processed as above. Cultures were plated onto 0.4% glucose (w/v) and 2×10^−4^% arabinose (w/v) and incubated overnight at 37 °C before imaging.

For producing Effector-Eco9 in a Δ*smpB* background, *E. coli* MG1655 containing the *smpB* deletion and containing pBAD33::Effector-Eco9 or a corresponding empty vector were grown to saturation and processed as above. Cultures were plated onto LB plates containing 0.2% arabinose (w/v) and incubated overnight at 37 °C before imaging.

For producing Effector-Eco9 in a Δ*smrB*, Δ*rnr*, Δ*arfA*, or Δ*arfB* background, *E. coli* MG1655 containing the indicated deletion allele and containing pBAD33::Effector-Eco9 or a corresponding empty vector were grown to saturation and processed as above. Cultures were plated onto 0.4% glucose (w/v) and 2×10^−2^% arabinose (w/v) and incubated overnight at 37 °C before imaging.

For producing Effector-Eco9-FLAG, *E. coli* MG1655 containing pBAD33::Effector-Eco9-FLAG were grown to saturation and processed as above. Cultures were plated onto 0.4% glucose (w/v) and 0.2% arabinose (w/v) and incubated overnight at 37 °C before imaging.

### Luminescence assay

*E. coli* MG1655 strains containing pBAD33::Effector-Eco9, pBAD33::*dam* and pBR322::Retron-Eco1, or the corresponding empty vector plasmids were cultured overnight in LB + 0.4% glucose (w/v) at 37 °C. Overnight cultures were diluted 1:100 in fresh LB (no glucose) and grown at 37 °C to an OD_600_ of approximately 0.3. Cells were diluted to OD_600_ of 0.2 in 15 mL LB + 0.2% arabinose (w/v) and grown at 37 °C for 30 minutes. Cells were then pelleted and resuspended to OD_600_ of 1.0 in 1X PBS. Samples were mixed with an equal volume of NAD/NADH-Glo detection reagent (Promega), and luminescence was measured on a Biotek plate reader with continuous shaking at 25 °C. Luminescence values 60 minutes after assay induction were reported as the mean and standard deviation of two biological replicates, each with three technical replicates.

### Incorporation assays

To produce Effector-Eco9, single colonies of *E. coli* MG1655 harboring pBAD33::Effector-Eco9 or the corresponding empty vector were grown overnight in LB + glucose + antibiotics. Overnight cultures were diluted to an OD_600_ of 0.05 in 25 mL M9 media containing 0.4% (w/v) glucose and grown at 37 °C to an OD_600_ of approximately 0.3. Cells were pelleted at 4,000 x *g* for 5 minutes at 4 °C, washed once with M9 media (no glucose), and then diluted to an OD_600_ of 0.1 in 10 mL M9 media (no glucose). Cultures were allowed to recover for 45 minutes at 37 °C before induction with 0.2% (w/v) arabinose (t = 0 minutes). At the indicated time points (0, 15, 30, 45, and 60 minutes), OD_600_ was measured, and a 250 µL aliquot of cells was transferred to a microcentrifuge tube containing [5,6-^3^H]-uridine (Revvity) (2-4 µCi/mL) for transcription measurements, [methyl-^3^H]-thymidine (Revvity) (20 µCi/mL) for DNA replication measurements, or EasyTag EXPRESS-^35^S Protein Labeling Mix (Revvity) (4 µCi/ mL) for translation measurements. Tubes were incubated at 37 °C for 2 minutes and quenched by adding nonradioactive uridine (1.67 mM), thymidine (1.67 mM), or cysteine and methionine (16.67 mM each), followed by an additional two-minute incubation. Samples were precipitated by adding ice-cold trichloroacetic acid (TCA) (10% w/v final concentration) and incubated on ice for at least 30 minutes. Precipitated material was collected on glass microfiber filters (Whatman, 1820-24) that were pre-wetted with 5% (w/v) TCA. Filters were washed sequentially with 35 volumes of 5% (w/v) TCA, 5 volumes of 100% ethanol, air dried, and transferred to scintillation vials containing scintillation fluid for measurements using a PerkinElmer scintillation counter. Counts per minute (CPM) were normalized to OD_600_ and presented relative to t = 0. Data represent the mean and standard deviation from two independent biological replicates.

### Yeast protein purification

Yeast strains CRB716 or CRB826 were grown to saturation overnight in YPD at 30 °C. Overnight cultures were used to inoculate 4 L YEP + 2% (w/v) raffinose at OD_600_ 0.2 and grown for ~15 hrs. When cultures reached OD_600_ ~0.7, 2% (w/v) galactose was added to induce protein expression for 2 hrs at 30 °C. Cells were harvested and washed sequentially with 0.1M PMSF, cell wash buffer (25 mM HEPES-KOH pH 7.6, 1M sorbitol), and lysis buffer (50 mM HEPES-KOH pH 7.6, 10% glycerol, 5 mM Mg(OAc)_2_, 1 mM EDTA, 1 mM EGTA, 250 mM KCl, 2 mM DTT, 0.2% NP-40, 2X cOmplete protease inhibitor tablets), before being resuspended in 10 mL lysis buffer + 1 mM PMSF and frozen dropwise in liquid nitrogen. Cells were lysed by grinding frozen cell pellets using a SPEX SamplePrep Freezer/Mill. After thawing, 15 mL Buffer H (50 mM HEPES-KOH pH 7.6, 10% glycerol, 5 mM Mg(OAc)_2_) containing 0.02% NP-40 and one cOmplete protease inhibitor tablet (Roche) was added to the cell lysate and centrifuged at 150,000 x *g* for 90 minutes in a SW32 Ti rotor (Beckman). The supernatant was filtered and mixed with 0.6 mL anti-FLAG affinity gel (Sigma) equilibrated with Buffer H. The lysate-resin mix was incubated for 4 hours at 4 °C with rotation. The resin was then washed with 5 volumes of Buffer H containing 1 mM EDTA, 1 mM EGTA, 250 mM KCl, and 2 mM DTT. Effector was eluted with 3 volumes of Buffer H containing 1 mM EDTA, 1 mM EGTA, 250 mM KCl, and 0.15 mg/mL 3XFLAG peptide. Elution fractions were concentrated with Amicon Ultra centrifugal filters with a 3 kDa MWCO, and protein concentration was measured by Bradford assay.

### tRNA purification

*E. coli* MG1655 strains containing pBAD33::Effector-Eco9, pBAD33::CapRel^SJ46^-N-term, or the corresponding empty vector plasmid were cultured overnight in LB + 0.4% glucose (w/v) at 37 °C. Overnight cultures were then diluted 1:50 in fresh LB (without glucose) and grown at 37 °C until OD_600_ of approximately 0.6 was reached. Toxin production was induced by adding 0.2% arabinose (w/v), and at indicated time points, 10 mL culture was harvested. Cells were pelleted by centrifugation at max speed for 10 minutes, the supernatant was discarded, and the pellet was resuspended in 1 mL TRIzol reagent (Invitrogen). Samples were flash frozen at −80 °C until RNA extraction.

TRIzol samples were thawed at room temperature, and 200 µL chloroform was added to each sample. Samples were vortexed for 30 seconds, then incubated on ice for 2 minutes. Tubes were briefly shaken to remix separated phases, then samples were spun at 5000 rpm for 15 minutes at 4 °C. 500 µL of the upper aqueous phase was transferred into a new microcentrifuge tube with 500 µL ice-cold isopropanol. Samples were vortexed for 15 seconds, then incubated at −20 °C for 1 hour. RNA was pelleted by centrifuging samples at max speed for 30 minutes. Pellets were washed three times with 70% ethanol, air dried, and resuspended in 90 µL DEPC water. For DNase treatment, 10 µL 10X TURBO DNase buffer (Invitrogen) and 2 uL TURBO DNase I (Invitrogen) was added to each sample. Samples were incubated at 37 °C for 20 minutes, another 2 µL TURBO DNase I was added, and samples were incubated at 37 °C for an additional 20 minutes. Samples were brought to 200 µL total volume with DEPC water, and 200 µL acid phenol:chloroform (125:24:1 phenol:chloroform:isoamyl alcohol, pH 4.5; Invitrogen) was added to each sample. Tubes were then vortexed for 1 minute and centrifuged at max speed for 10 minutes at 4 °C. The aqueous layer was transferred to a new microcentrifuge tube, and 0.1 volumes 3M NaOAc, 3 volumes 100% ethanol, and 0.01 volumes GlycoBlue (Invitrogen) were added to each sample. RNA was precipitated at −20 °C for at least 1 hour and then pelleted by centrifuging tubes at max speed for 30 minutes at 4 °C. Pellets were washed twice with 70% ethanol, air dried, and resuspended in DEPC water.

Following RNA extraction, tRNA species were further purified from acrylamide gels using previously described crush-and-soak gel extraction methods^52,61^. Briefly, approximately 25 µg of total cellular RNA isolated 0 or 60 minutes after toxin induction was resolved across several wells of an 8% TBE-Urea acrylamide gel. After running, the gel was then stained with SYBR Gold (Invitrogen), and all RNA species below the 5S rRNA were cut out into 1 cm x 1 cm squares and combined into a single tube. Gel fragments were weighed, and 2 volumes (v/w) of elution solution (500 mM NH_4_OAc, 1 mM EDTA-KOH pH 8.0) was added. Samples were incubated for 2 hours at room temperature, followed by an overnight incubation at 4 °C. The following day, samples were spun at max speed for 5 minutes, and the supernatant was collected. To precipitate the RNA, 0.1 volumes of 2M NaOAc pH 5.2 and 3 volumes of ethanol were added to the supernatant, and the mixture was incubated at −20 °C for 1 hour. RNA was pelleted by centrifuging the sample at 5,000 x *g* at 4 °C for 30 minutes. The RNA pellet was washed twice with cold 75% ethanol, air dried, and resuspended in DEPC water.

### *In vitro* transcription/translation reactions

#### ΔtRNA reactions

*In vitro* transcription/translation reactions were performed using the PURExpress reconstituted protein synthesis system (NEB) lacking tRNAs (ΔtRNA kit). Assays were performed following the manufacturer’s instruction, scaled down to 11 µL (ΔtRNA experiments) reactions containing 250 ng template DNA. To monitor protein production by Western blot, the template DNA provided by the manufacturer (pT7::*folA*) was modified to include a C-terminal FLAG tag (pT7::*folA*-FLAG). 600 ng gel extracted tRNA was added to each 11 µL reaction. Reactions were incubated at 37 °C for 2 hours and then mixed 1:1 with 2X Laemmli sample buffer (Bio-Rad) containing 10% (v/v) 2-mercaptoethanol. Samples were boiled at 95 °C for 10 minutes, and 10 µL each reaction was loaded into a 4-20% SDS-PAGE gel (Bio-Rad). The gel was run at 100V for 10 minutes and then 150V for 40 minutes. Total protein was wet transferred to a 0.2 µM PVDF membrane at 100V for 60 minutes. Membranes were blocked in Tris-buffered saline with 0.1% Tween-20 (TBST) and 5% non-fat milk for 30-60 minutes at room temperature and then incubated with primary monoclonal antibody (1:1,000 rabbit anti-FLAG, Cell Signaling Technologies) overnight at 4 °C. Membranes were washed with TBST and incubated with HRP-conjugated goat anti-rabbit IgG (Invitrogen) in TBST + 5% milk for 60 minutes at room temperature. Membranes were washed again and incubated with SuperSignal West Femto Maximum Sensitivity Substrate (Thermo Scientific) before exposure on a Bio-Rad ChemiDoc MP imager. Membranes were stained with Coomassie as a loading control. Images shown are representative of two replicates from one tRNA purification.

#### Reactions containing purified effector

*In vitro* transcription/ translation reactions were performed using the PURExpress reconstituted protein synthesis system following the manufacturer’s instruction. Reactions were scaled down to 12.5 µL and contained 125 ng template DNA. 1 mM NAD^+^ was added to reactions where indicated. Purified effector was added at the following concentrations: 2 µM (**Figures 2A, 2C and 3A**), 0.06 µM – 2 µM (**Figure 3B**), and 1.4 µM (**Figures 3C, 3D, S2G, and 4H**). Reactions were incubated at 37 °C for 2 hours, unless otherwise indicated, and then mixed 1:1 with 2X Laemmli sample buffer (Bio-Rad). 5-10 µL each reaction was loaded into a 4-20% SDS-PAGE gel (Bio-Rad, **Figure 2A**) or a 12% NuPAGE Bis-Tris gel (Invitrogen, all other figures). SDS-PAGE gels were run at 100V for 10 minutes then 150V for 40 minutes, and Bis-Tris gels were run at 120V for 120 minutes. SDS-PAGE gels were stained in Coomassie before imaging. Total protein from Bis-Tris gels was wet transferred to a 0.45 µM PVDF membrane at 70V for 120 minutes. Membranes were blocked in Tris-buffered saline (TBS) and 3% BSA for 30 minutes at room temperature and then incubated with TBS with 0.05% Tween-20 (TBST), 1% BSA, and primary monoclonal antibody (1:1000 rabbit anti-ADPr, rabbit anti-FLAG, mouse anti-HA, or rabbit anti-Myc (Cell Signaling Technologies)) overnight at 4 °C. Membranes were washed with TBST and incubated with HRP-conjugated goat anti-rabbit or anti-mouse IgG (Invitrogen) in TBST + 1% BSA for 1 hour at room temperature. Membranes were washed again and incubated with SuperSignal West Femto Maximum Sensitivity Substrate (Thermo Scientific) before exposure on a Bio-Rad ChemiDoc MP imager. Membranes were stained with Coomassie as a loading control. Images shown are representative of two replicates.

For *in vitro* reactions loaded onto sucrose gradients, reactions were prepared in 25 µL volume with 250 ng pT7::HA-DHFR-TGA DNA template. Reactions were incubated at 37 °C for 20 minutes, and then immediately loaded onto a 10-55% (w/v) sucrose gradient (20 mM Tris-HCl pH 7.6, 50 mM MgCl_2_, 100 mM NH_4_Cl, 1 mM DTT, 0.5 mM EDTA, 10% or 55% (w/v) sucrose) generated using a Gradient Master (BioComp). Gradients were centrifuged at 35,000 rpm for 2.5 hours at 4 °C in a SW41-Ti rotor and fractioned using the Piston Gradient Fractionator (BioComp) with continuous A_260_ measurements. Plotted A_260_ values were smoothed using a Savitsky-Golay filter with a window length of 100 and a polyorder value of 3.

For *in vitro* reactions treated with nuclease, 12.5 U benzonase (Millipore) was added to the reaction after addition of Laemmli buffer. Samples were incubated at 37 °C for 30 minutes before being loaded into a Bis-Tris gel, as previously described.

### RNA dot blots

*E. coli* MG1655 strains containing pBAD33::Effector-Eco9 or the corresponding empty vector plasmid were cultured overnight in LB + 0.4% glucose (w/v) at 37 °C. Overnight cultures were then diluted 1:100 in fresh LB (without glucose) and grown at 37 °C until OD ~0.2 was reached. Toxin production was induced by adding 0.2% arabinose (w/v), and at indicated time points, 900 µL cells were added to 100 µL stop solution (95% ethanol (v/v) and 5% phenol solution saturated with citrate (v/v)). Cells were pelleted at 13,000 x *g* for 30 seconds, the supernatant removed, and flash frozen at −80 °C until sample collection was complete. 400 µL TRIzol (Invitrogen) heated to 65 °C was then added to frozen cells, and samples were incubated at 65 °C for 10 minutes at 2000 rpm on a thermomixer. Samples were then flash frozen at −80 °C for 10 minutes, thawed at room temperature for 5 minutes, and then pelleted for 5 minutes at max. speed at 4 °C. The supernatant was then combined 1:1 with ethanol and added to a Direct-Zol RNA Miniprep (Zymo) spin column. Samples were centrifuged at 13,000 x *g* for 30 seconds, and the flow through discarded. The column was washed twice with RNA PreWash Buffer and once with RNA Wash Buffer, as described in the Direct-Zol kit. RNA was then eluted with RNase-free water. DNA was then depleted from RNA samples using TURBO DNase (Invitrogen), as described in the ‘tRNA purification’ section.

1 µg RNA was then spotted onto Amersham Hybond-N+ membrane (Cytiva), saturated with 1X TBE. RNA was crosslinked to the membrane using a Stratagene UV Stratalinker 1800 set. The membrane was washed once in TBST and then blocked with TBST + 5% milk (w/v) for 1 hour. The membrane was then incubated with primary antibody (rabbit anti-ADPr, Cell Signaling Technologies) at a 1:1000 dilution in TBST + 5% milk (v/v) overnight. The membrane was then washed three times with TBST for 5 minutes each and then incubated with secondary antibody (HRP-conjugated goat anti-rabbit or anti-mouse IgG, Invitrogen) for 1 hour. The membrane was washed again three times for 5 minutes each with TBST and incubated with SuperSignal West Femto Maximum Sensitivity Substrate (Thermo Scientific) before exposure on a Bio-Rad ChemiDoc MP imager. Membranes were stained with methylene blue solution (0.04% methylene blue (w/v), 0.5M NaOAc) as a loading control.

### Immunoblots to detect ADPr from cell lysates

To detect ADPr modifications on proteins from cell lysates, *E. coli* MG1655 strains containing pBAD33::Effector-Eco9, pBAD33::Effector-Eco9(E108A), or the corresponding empty vector were cultured overnight in LB + 0.4% glucose (w/v) at 37 °C. Overnight cultures were diluted 1:100 in fresh LB (without glucose) and grown at 37 °C until OD ~0.3 was reached. Cultures were diluted to OD_600_ 0.2 in LB, and toxin production was induced by adding 0.2% arabinose (w/v). At indicated time points, 10 mL culture was harvested and pelleted at 4,000 x *g* for 10 minutes. Pellets were resuspended in 50 µL lysis buffer (50 mM Tris-HCl pH 8.0, 150 mM NaCl, 2% SDS, 1 mM MgCl_2_, cOmplete Protease Inhibitor Cocktail (Roche)) for every 1.0 OD unit. Lysates were treated with 12.5 U benzonase (Millipore) at room temperature for 10 minutes. Lysates were diluted 1:3 in Laemmli buffer and samples were then pelleted at 16,000 x *g* for 5 minutes, saving the supernatant. 10 µL each sample was loaded into a 12% NuPAGE Bis-Tris gel. Gel electrophoresis and immunoblotting was then performed as described in the ‘*in vitro* transcription/translation reactions containing purified effector’ section.

To detect ADPr modification in cells co-producing Effector-Eco9 and RF1 or RF2 (or 3XFLAG-tagged variants), *E. coli* MG1655 strains containing the indicated plasmids were cultured overnight in LB + 0.4% glucose (w/v) at 37 °C. Overnight cultures were diluted 1:100 in fresh LB (without glucose) and 20 µM vanillic acid and grown at 37 °C until OD ~0.3 was reached. Cultures were diluted to OD_600_ 0.2 in LB + 20 µM vanillic acid, and toxin production was induced by adding 0.2% arabinose (w/v). Cell lysis, gel electrophoresis, and immunoblotting were then performed as described for above samples. For anti-EF-Tu blots, mouse anti-eEF1A1/EF-Tu antibody (Abcam) was used as the primary antibody in a 1:1000 dilution.

### *In vitro* RF1/RF2 ADP-ribosylation reactions

~400 ug/mL purified RF1 or RF2 from NEB PURExpress ΔRF1/2/3 kits were combined with 2 µM purified effector and 1 mM NAD^+^ in buffer containing 50 mM HEPES-KOH pH 7.6, 10% glycerol, 5 mM Mg(OAc)_2_, 1 mM EDTA, 1 mM EGTA, and 250 mM KCl. Samples were incubated at 37 °C, and at indicated time points, the reaction mixture was combined with 1X Laemmli buffer and then loaded onto 12% NuPAGE Bis-Tris gels (Invitrogen). Gel electrophoresis and immunoblotting were then performed as described in the ‘*in vitro* transcription/translation reactions containing purified effector’ section.

### Co-immunoprecipitation and LC-MS/MS

Overnight cultures of pBAD33::Effector-Eco9 E108A or pBAD33::Effector-Eco9 E108A-FLAG (both also containing an empty pVan control plasmid) were diluted in 75 mL LB + 0.2% (w/v) arabinose to OD_600_ 0.05 and grown at 37 °C to an OD_600_ of 0.3. 100 µM vanillic acid was added to each culture, and cultures were grown at 37 °C for an additional hour. 50 mL of each sample was then pelleted by centrifugation at 4,000 x *g* for 10 minutes at 4 °C. The supernatant was discarded, and cell pellets were resuspended in 900 µL lysis buffer (25 mM Tris-HCl, 150 mM NaCl, 1 mM EDTA, 5% glycerol, 1% Triton X-100) supplemented with 1 µL/mL ReadyLyse Lysozyme (Fisher Scientific), 1 µL/mL benzonase (Millipore), and cOmplete Mini Protease Inhibitor Cocktail (Roche) and then flash frozen at −80 °C. Two additional freeze-thaw cycles in liquid nitrogen were used to ensure complete cell lysis. Additional lysis buffer was then added to normalize sample concentrations by OD_600_ taken at the time of cell harvesting. Samples were spun at 20,000 x *g* for 10 minutes at 4 °C to pellet debris. For each sample, 50 µL Pierce anti-DYKDDDDK magnetic agarose beads (Thermo Scientific) were combined with 450 µL wash buffer (25 mM Tris-HCl, 150 mM NaCl, 1 mM EDTA, 5% glycerol, cOmplete mini Protease Inhibitor Cocktail (Roche)) and collected using a magnetic rack. Beads were then washed twice with 500 µL wash buffer, then incubated with 850 µL sample for 1 hour at 4 °C on an end-to-end rotor. After incubation, beads were washed twice in 1X PBS and once with MilliQ H_2_O. To validate the IP before LC-MS/MS (**Figure S3A**), a small aliquot of beads was combined with 1X Laemmli buffer and boiled at 95 °C for 10 minutes. Samples were then run on an SDS-PAGE gel and stained with Coomassie as described in the ‘*in vitro* transcription/translation ΔtRNA’ section. The remaining beads were used for downstream LC-MS/MS.

On-bead reduction, trypsin digest, and LC-MS/MS were done by the MIT Biopolymers and Proteomics Core as previously described^8,20^. In brief, proteins were reduced for 1 hour at 56 °C with 10 mM dithiothreitol (Sigma) and then alkylated for 1 hour at 25 °C in the dark with 20 mM iodoacetamide (Sigma). Proteins were digested with modified trypsin (Promega) overnight in 100 mM ammonium bicarbonate pH 8. Formic acid (99.9%, Sigma) was added to stop trypsin digest at a final concentration of 5%. Digested peptides were desalted using Pierce Peptide Desalting Spin Columns (Thermo) then vacuum centrifuged.

Peptides were separated on a PepMap RSLC C18 column (Thermo) over a 90 min gradient by reverse phase HPLC (Thermo Ultimate 3000) before nano-electrospray with an Orbitrap Exploris 480 mass spectrometer (Thermo). The mass spectrometer was run in a data-dependent mode. Full scan parameters were resolution of 120,000 across 375– 1600 *m*/*z* and maximum IT 25 ms. This was followed by MS/ MS for as many precursor ions in a two second cycle with a resolution of 30,000, dynamic exclusion of 20 s, and a NCE of 28. Detected peptides were mapped to *E. coli* MG1655 using Sequest HT in Proteome Discoverer (Thermo). The ratio of spectral counts between the FLAG-tagged pulldown and untagged pulldown at each time point were used to generate figure data, with a pseudocount added to each count. Proteins with less than 50 spectral counts in the FLAG condition were excluded from analysis.

### Sucrose gradients following effector induction

*E. coli* MG1655 strains containing pBAD33::Effector-Eco9 or the corresponding empty vector plasmid were cultured overnight in LB + 0.4% glucose (w/v) at 37 °C. Overnight cultures were diluted 1:50 in fresh LB (without glucose) and grown at 37 °C until OD_600_ ~0.3 was reached. Effector production was induced by adding 0.2% arabinose (w/v), and at indicated time points, 50 mL culture was harvested. Cells were pelleted by spinning the culture at 7800g for 10 minutes at 4 °C, the supernatant was discarded, and the pellet was flash frozen at −80 °C. Cell pellets were then thawed on ice and resuspended 1:2 (w/v) in homogenization buffer (20 mM Tris-HCl pH 7.6, 50 mM Mg(OAc)_2_, 100 mM NH_4_Cl, 1 mM DTT, 0.5 mM EDTA). Resuspended pellets were lysed by sonication (6 x 15 second pulses at an amplitude of 25%), and lysates were cleared by centrifugation at 30,000 x *g* for 30 minutes at 4 °C. Cleared lysate corresponding to 0.3 mg total RNA (as measured by Nanodrop) was layered over a 10-55% (w/v) sucrose gradient. Gradients were centrifuged, fractionated, and plotted with smoothing as described previously.

### Ribosome profiling

Ribosome profiling was performed as previously described^62^ with minor modifications. Briefly, overnight cultures of *E. coli* MG1655 containing pBAD33::Effector-Eco9 or no additional plasmids were diluted to OD_600_ 0.005 in 200 mL LB (no glucose) and grow at 37 °C until OD_600_ of ~0.3 was reached. Production of Effector-Eco9 in cultures containing pBAD33::Effector-Eco9 was then induced by adding 0.2% (w/v) arabinose, and at indicated time points, cells were harvested by filtration over a 90 mm 0.22 µM PVDF filter (Millipore) attached to a vacuum flask. Immediately following filtration, cells were scraped off the filter and frozen in liquid nitrogen. For cell lysis, lysis buffer (20 mM Tris pH 8.0, 100 mM NH_4_Cl, 10 mM MgCl_2_, 0.4% Triton X-100, 0.1% NP-40, 1 mM chloramphenicol, 100 U/mL RNase-free DNase (Thermo Scientific)) was first flash frozen in individual droplets by pipetting into liquid nitrogen. Frozen lysis buffer was combined with frozen cells in stainless steel grinding jars (Qiagen) that were pre-cooled in liquid nitrogen. Cells were then lysed on a TissueLyser II (Qiagen) three times at 15 Hz for 3 minutes, cooling jars in between each interval. Lysate was then thawed and centrifuged at 20,000 rpm for 10 minutes at 4 °C to pellet debris. Supernatant was transferred to a new tube, and the approximate concentration of RNA in the sample was determined by measuring absorbance at 260 nm. Then, supernatant containing 0.5 mg RNA was combined with 750 U Nuclease S7 (MNase) (Millipore Sigma), 100 U SUPERase•In (Invitrogen), and 5 mM CaCl_2_ in 300 µL total volume, adding additional lysis buffer if necessary. Samples were incubated at 25 °C for 1 hour, and then quenched by adding 5 mM EGTA. Digested sample was then layered over a 10-55% (w/v) sucrose gradient in buffer containing 20 mM Tris pH 8.0, 100 mM NH_4_Cl, 10 mM MgCl_2_, and 1 mM chloramphenicol. Gradients were centrifuged at 35,000 rpm for 2.5 hours in a SW41-Ti rotor and fractioned using a Piston Gradient Fractionator (BioComp) with continuous A260 measurements. Monosome fractions were collected, and flash frozen at −80 °C until RNA extraction.

To isolate RNA from monosome fractions, fractions were first thawed, and 1% SDS was added to each sample. Acid phenol chloroform pH 4.5 (ThermoFisher) was then added to each sample 1:1 (v/v), and samples were incubated at 65 °C for 5 minutes at 1,400 rpm in a thermomixer. Samples were chilled on ice for 5 minutes and then centrifuged at 20,000 x *g* for 2 minutes at room temperature. The top aqueous layer was transferred to a new tube, and acid phenol chloroform pH 4.5 was again added to the sample in a 1:1 (v/v) ratio. Samples were incubated at room temperature for 5 minutes, vortexing occasionally. Samples were centrifuged and the top aqueous layer extracted as in the previous step. The aqueous layer was then combined with chloroform in a 1:1 (v/v) ratio and vortexed for 30 seconds at room temperature. The top aqueous layer was transferred to a new tube, and RNA was precipitated by adding 0.1 volumes 3 M NaOAc, 1 volume isopropanol, and 2 µL GlycoBlue (Invitrogen). Samples were incubated at −80 °C for 30 minutes before being centrifuged at 20,000 x *g* for 20 minutes at 4 °C to pellet RNA. Pellets were washed once in 80% ethanol, air dried, and resuspended in 10 mM Tris pH 7.

Approximately 20 µg RNA was run across two wells in a 15% TBE-Urea PAGE gel and stained with SYBR Gold (Invitrogen). Bands between 15 and 45 nucleotides were extracted using a clean razor blade. Gel pieces were placed in a 0.5 mL tube with a hole in the bottom, placed within a 2 mL tube. Gel pieces were then pulverized by centrifugation at 20,000 x *g* for 3 minutes at room temperature. 500 µL 0.5 M NH_4_Cl was added to crushed gel pieces, and the mixtures was incubated at room temperature for 1 hour (stationary) and then overnight at 4 °C on an end-to-end rotor. The gel mixture was then transferred to a Spin-X cellulose acetate column (Corning) and centrifuged at 20,000 x *g* for 3 minutes. The eluate was transferred to a new tube, brought to 500 µL total volume with 10 mM Tris pH 7, and combined with 0.1 volumes NaOAc, 1 volume isopropanol, and 2 µL GlycoBlue (Invitrogen). RNA was precipitated for at least 30 minutes at −80 °C and then centrifuged at 20,000 x *g* for 30 minutes at 4 °C. The RNA pellet was washed with 80% ethanol, air dried, and resuspended in 10 mM Tris pH 7.

RNA was dephosphorylated by combining size selected RNA, 1X T4 PNK buffer (New England Biolabs), 1 U/µL Superase•In (Invitrogen), and 1 U/µL T4 PNK (New England Biolabs). The reaction was incubated at 37 °C for 1 hour and then heat inactivated 75 °C for 10 minutes. The reaction was then brought to 500 µL total volume with additional 10 mM Tris pH 7 and precipitated as described previously. RNA was then ligated to linker-1 (CB204) by combining dephosphorylated RNA with 25% PEG 8000, 1X T4 RNA ligase 2 buffer (New England Biolabs), 5 µM linker-1, and 10 U/µL T4 RNA ligase 2 truncated K227Q. The ligation reaction was incubated at 25 °C for 2.5 hours before the total volume was brought to 500 µL with 10 mM Tris pH 7, and RNA was precipitated as described previously. Ligated RNA was then run on a 10% TBE-Urea gel, and bands between 35 and 65 nucleotides were gel extracted and precipitated as described previously.

Ligated RNA was then reverse transcribed by combining precipitated RNA, 0.5 mM dNTPs (Invitrogen), 1.25 µM oCJ485 (CB205), 1X First Strand Buffer, 1 U/µL Superase•In (Invitrogen), 5 mM DTT, and 10 U/µL SuperScript III (Invitrogen). The reverse transcription reaction was incubated at 50 °C for 30 minutes before adding 100 mM NaOH and heating at 95 °C for 15 minutes to hydrolyze RNA. The reaction was then directly loaded into a 10% TBE-Urea gel, and bands between 90 and 120 nucleotides were gel extracted. Gel pieces were pulverized and incubated in 0.5 M NH_4_Cl as described previously. The gel mixture was then transferred to a Spin-X cellulose acetate column (Corning) and centrifuged at 20,000 x *g* for 3 minutes. The eluate was transferred to a new tube, brought to 500 µL total volume with 10 mM Tris pH 8, and combined with 150 mM NaCl, 500 µM EDTA, 1 volume isopropanol, and 2 µL GlycoBlue (Invitrogen). DNA was precipitated at −80 °C for at least 30 minutes and then centrifuged at 20,000 x *g* for 30 minutes at 4 °C. The DNA pellet was washed with 80% ethanol, air dried, and resuspended in 10 mM Tris pH 8.

cDNA was then circularized by combining cDNA, 50 µM ATP, 1X CircLigase Buffer (Biosearch Technologies), 2.5 mM MnCl_2_, and 5 U/µL CircLigase (Biosearch Technologies). The reaction was incubated at 60 °C for 1 hour before an additional 100 U CircLigase was added to the reaction and incubated again at 60 °C for 1 hour. The reaction was inactivated by heating at 80 °C for 10 minutes and returned to ice. rRNA subtraction was then performed by combining 5 µL cDNA with rRNA subtraction oligo mix (7.7 µM o1055 (CB697), 0.4 µM o1056 (CB698), 1.7 µM o1057 (CB699), and 0.2 µM o1058 (CB700)) and 2X SSC. The reaction was incubated at 98 °C for 75 seconds before ramping down to 37 °C over one hour, followed by a final incubation at 37 °C for 20 minutes. MyOne Streptavidin C1 Dynabeads (Invitrogen) were then washed three times with 1 volume of 1X B&W buffer (5 mM Tris pH 7.5, 0.5 mM EDTA, 1 M NaCl, 0.01% Tween) and then resuspended in 0.4 volumes 2X B&W buffer (10 mM Tris pH 7.5, 1 mM EDTA, 2M NaCl, 0.01% Tween). Washed beads were incubated at 37 °C before an equal volume of beads was added to the hybridization reaction and incubated at 37 °C for 15 minutes. The eluate was transferred to a new tube, brought to 500 µL total volume with 10 mM Tris pH 8, and precipitated as described previously for DNA samples.

Final sequencing libraries were then amplified by combining rRNA subtracted cDNA, 1X HF buffer (New England Biolabs), 0.2 mM dNTPs, 0.5 µM o231 (CB206), 0.5 µM indexing primer (CB705, CB706, or CB707), and 0.02 U/µL HF Phusion (New England Biolabs). Following an initial melting at 98 °C for 30 seconds, reactions were subjected to 12-24 cycles of denaturation at 98 °C for 10 seconds, annealing at 60 °C for 10 seconds, and extension at 72 °C for 5 seconds. DNA was then run on a 8% TBE-PAGE gel, and bands approximately 175 base pairs were excised, gel extracted, and precipitated as described previously for DNA samples. Single-end sequencing was performed on an Illumina NextSeq at the MIT BioMicroCenter.

### Ribosome profiling sequencing analysis

Single-end ribosome profiling reads were processed using a custom Python pipeline (v3.12.12) created with assistance from ChatGPT. First, 3’ adapter sequences (CTGTAGGCACCATCAAT) were removed using Cutadapt (v5.1). Reads were then filtered at a Phred quality score of 30 (-q 30) and reads shorter than 15 nt were discarded (-m 15). Reads were then again filtered to retain fragments 20-42 nts in length. Bowtie v1.3.1 was used to remove reads mapping to annotated rRNA and tRNA sequences, allowing up to two mismatches per alignment (-v 2) and excluding reads with more than one alignment (-m 1). Remaining reads were aligned to the phiX174 reference sequence (NC_001422) using the same Bowtie parameters, and mapped reads were discarded. Reads were then aligned to the MG1655 genome (U00096.3) using Bowtie2 v2.5.5 in single-end mode with the --very-sensitive preset. SAM alignments were converted BAM format, coordinate-sorted, and indexed using SAMtools v1.23.

For metagene analyses, coding sequences (CDSs) were identified from the MG1655 GFF3 annotation, and CDSs shorter than 100 nt were excluded from analysis. For each aligned read, the ribosomal P-site was estimated from the 3’ end of the read using a 15 nt offset. Each CDS was divided into 50 equal length bins spanning the entire coding region. P-sites were assigned to bins according to their relative position within the CDS, and the number of P-sites in each bin was recorded for each CDS. Counts for each CDS were then normalized by the total number of P-sites assigned to that CDS. Normalized profiles were then averaged across all qualifying CDSs to generate the metagene profile for each sample. For flanking regions, P-sites were counted for each nucleotide ±15 nt of the corresponding CDS. Counts at each nucleotide position were then normalized by the total number of P-site counts for the corresponding CDS, and normalized profiles were averaged across flanking regions of all qualifying CDSs.

### CryoEM sample preparation

CryoPRISM lysates were generated as detailed previously^27^. Briefly, *E. coli* strain MG1655 bearing an inducible effector plasmid (pCB48) or empty vector control plasmid (pCB120) was grown with aeration at 37 °C in LB medium supplemented with chloramphenicol and glucose (0.4% w/v). Cells grown to a density of OD_600_ ~0.25 were induced by addition of arabinose (0.2% w/v) for 30 minutes, before harvesting by quenching on ice and pelleting by centrifugation at 4,000 *g* for 15 min in a Beckman JLA-8.100 rotor. Final cell densities reached OD_600_ ~0.4 and OD_600_ ~0.5 for the + effector and – effector samples, respectively. Cell pellets were scraped and flash frozen in liquid nitrogen, followed by cryogenic cell lysis using a Retsch MM-400 Mixer Mill with a 5 mL stainless steel grinding jar and 10 mm stainless steel grinding balls with five milling cycles at 30 Hz for 90 s. Frozen lysate was then resuspended in ice-cold buffer A (20 mM Tris-HCl, 100 mM NH_4_Cl, 10 mM MgCl_2_, 0.5 mM EDTA, 6 mM β-mercaptoethanol, pH 7.5) and clarified by centrifugation (21,000 x *g*, 20 min, 4 °C) before addition of 2 µL DNase I (33 U/mL of lysate) and 1X DNase I Reaction Buffer (New England Biolabs #M0303L and #B0303S, respectively). DNase-treated lysate was further clarified by centrifugation (21,000 x *g*, 10 min, 4 °C).

*In vitro* transcription/translation reactions for cryoEM were performed using the PURExpress reconstituted protein synthesis system (NEB) following the manufacturer’s instruction. Reactions were prepared in 25 µL total volume containing 250 ng template DNA (pCB362), 1 mM NAD^+^, and 1.4 µM purified effector. Samples were incubated at 37 °C for 20 minutes before being placed on iced and loaded onto grids.

### CryoEM data collection and pre-processing

200-mesh Quantifoil R2/1 copper grids were coated with a graphene monolayer^27^ and rendered hydrophilic with a UV/ozone treatment 10 min using a Bioforce PC440 UV/ ozone cleaner. Sample (4 µL) was applied and plunge-frozen in liquid ethane using an FEI Vitrobot Mk IV instrument with a 4 s blot time and blot force of −2 at 10 °C and 95% relative humidity. CryoPRISM datasets were collected with EPU aberration-free image shift (AFIS) on a Titan Krios G3i operated at 300 kV using a Gatan K3 detector equipped with a BioQuantum energy filter. CryoPRISM datasets were collected with a total dose of 45.1 e−/Å fractionated across 45 frames; defocus ranged from −0.4 to −2.4 µm in steps of −0.2 µm; IVTT datasets were collected with a total dose of 50.8 e−/Å fractionated across 50 frames; defocus ranged from −0.2 to −2.2 µm. All datasets were collected at a magnification of 81,000X and a nominal pixel size of 1.06 Å.

Data for lysate and IVTT samples pre-processing was performed in cryoSPARC^31^ v4.5.3 and v5.0.6 as depicted (**Figure S4 - S11**) using default parameters for listed jobs, unless otherwise specified. Briefly, micrographs Patch Motion Correction and Patch CTF estimation preceded micrograph curation and particle picking (Blob Picker). Particle extraction (box size 440 pixels) produced 3,009,936 particles (+ effector cryoPRISM), 2,078,556 particles (− effector cryoPRISM), 4,385,360 particles (+ effector IVTT), and 3,491,027 (− effector IVTT). Particles were sorted via heterogenous refinement using six input volumes: a 70S ribosome, 30S ribosomal subunit, and 50S ribosomal subunit, and three “junk” volumes. Relevant classes were further filtered via multiclass *ab-initio* reconstruction, with particles sorting into “rejected” output classes excluded from each subsequent round of *ab-initio* reconstruction.

### CryoDRGN training and occupancy analyses

The 70S, 30S, and 50S particle stacks (downsampled to box size 128 pixels; 3.64 Å/pixel) were further analyzed for heterogeneity using cryoDRGN v2.2^28,29^ using estimated CTF parameters and particle poses from homogeneous reconstruction. CryoDRGN hyperparameters, MAVEn “*k*-means” analysis, and MAVEn “on-the-fly” analyses was performed as described previously^27^.

### Structural analysis of 70S particles and disomes

CryoDRGN-based filtering, guided by expert inspection of *k*-means centroid volumes for the – and + effector datasets, resulted in 609,658 and 209,050 70S particles, respectively. An additional 24,233 particles were removed from the + effector dataset through iterative 3D classification in cryoSPARC, resulting in 585,425 particles (**Figure 6A**), whereas 40,414 particles found to bear RF2 were removed from the − effector dataset, resulting in 168,636 particles. CryoDRGN models were trained on 70S particles stacks, with 1,000 volumes sampled by cryoDRGN from *k*-means cluster centers used for subsequent MAVEn “*k*-means” and voxel PCA (vPCA) analyses, which were performed as described previously^27^ at a box size of 128 pixels.

To separate 70S rotated and non-rotated particles, vPCA was performed using a mask designed to capture 30S body rotation relative to the 50S (**Figure S6A**), as described^27^. Thresholds for distinguishing rotational states are listed in **Table S1**. Particles in the rotated state with tRNAs in the hybrid positions were further separated based on the presence or absence of A/P tRNA using 3D classification on cryoSPARC (**Figure S6A**), resulting in 11,109 (+ effector) and 4,112 (− effector) particles with A/P and P/E tRNA and 3,202 particles with P/E tRNA and an empty A-site found exclusively in the – effector dataset. The remaining particles were analyzed by MAVEn “*k*-means” using masks for A/A, P/P, E/E-site tRNA in the non-rotated state, and a mask covering the GTPase-associated center (GAC) (**Figure S6A**), with occupancy thresholds defining 70S translational states (**Figure 6A**) listed in **Tables S2-3**. Particles comprising each sub-class were exported to cryoSPARC and used for homogeneous refinement jobs (**Figure 6A-B**). For the (+) effector dataset, the particles corresponding to the P/P, E/E-site tRNA-bound state were further classified using a mask around the nascent chain exit tunnel (NPET) (**Figure S6B**) to discriminate between particles with exit tunnel density and those without (**Figure 6B**).

Notably, cryoDRGN analysis of 70S particles produced a highly featured latent space, with disomes forming a distinct cluster (**Figure S6C**). These particle locations were imported to cryoSPARC, re-extracted at a larger box size (686 pixels, downsampled to 256 pixels) and used for *ab-initio* reconstruction followed by homogeneous refinement, yielding a disome map. Masks were generated for the leading and queueing ribosomes, and local refinement was applied to improve resolution of each disome component. The resulting local refinements were combined into a composite disome map using the “vop add” feature in ChimeraX (**Figure 6E**).

### CryoPRISM data analysis of 50S particles

The 50S particles containing density located near the P-site tRNA binding site (38,436 particles) or near the peptidyl transferase center (PTC) (35,377 particles) were identified within the + effector dataset (81,272) by inspection of cryoDRGN-generated volumes sampled from 50 *k*-means centroid locations. Associated particles were then imported to cryoSPARC for homogenous refinements. Aberrant 50S termination complexes 1 and 2 were identified via iterative 3D classification (cryoSPARC) using a large mask spanning the P-site tRNA binding site (**Figure S6D**), resulting in a particle stack with density near P-site tRNA and Hsp15, and a particles stack with only density near P-site tRNA. Aberrant 50S termination complex 3 was identified from particles with density near the PTC using a 2-class 3D classification job (cryoSPARC) with a mask spanning the PTC (**Figure S6D**). This yielded a class with density identified as HflX based on *de novo* atomic models built by ModelAngelo^63^ v1.0 using the build_no_seq function and subsequently assigned to the *E. coli* (K12) proteome (Uniprot ID UP00625) using probability-based scoring.

### CryoPRISM data analysis of 30S particles

The 30S particle stacks from the − effector dataset (49,988 particles) dataset were filtered by magnitude of the latent variable to remove particles producing poor reconstructions, resulting in 41,571 remaining particles. The 65,277 (+ effector) and 41,571 (− effector) particles (**Figure 6A**) were subjected to MAVEn “on-the-fly” analysis using masks for IF1, IF2, IF3, and tRNA (**Figure S6E**). Thresholds for distinguishing different 30S states are listed in **Tables S4-5**. Particles comprising each sub-class were exported to cryoSPARC and density maps were produced through homogeneous refinements (**Figure 6D**).

### IVTT cryoEM data analysis

The 70S particle stack from the + effector (384,754 particles) and – effector datasets (388,157 particles) were filtered by manual inspection of cryoDRGN-generated volumes sampled from *k*-means (*k*=1000 centroid locations to remove clusters of poorly resolved particles, resulting in 343,333 (+ effector) and 344,944 (− effector) particles. Ribosomes were first classified into rotated and non-rotated states using vPCA as described above, resulting in 50,821 (+ effector) and 57,904 (− effector) particles in the rotated state with tRNAs in the hybrid A/P and P/E states. Remaining particles were analyzed using MAVEn “*k*-means” analyses with masks for A/A, P/P, E/E-site tRNA and the GAC center as described above, with thresholds listed in **Table S6**. Particles classified as ribosomes with E/E, P/P, A/A tRNA and E/E, P/P, and an empty A-site for both datasets were exported to cryoSPARC for homogeneous refinements. Particles from each refinement were used for an additional round of cryoDRGN training and MAVEn analysis using 1,000 volumes sampled from *k*-means cluster centers from the latent space. Masks corresponding to adjacent ribosomes in the leading and queuing positions are used to query particles (**Figure S6F**). Maps with adjacent density for two adjacent ribosomes were classified as “polysomes,” with no adjacent density were classified as “monosomes,” and with adjacent density in only one position were classified as either “leading” or “queuing” based on the location of the adjacent density **Table S6**. All particles were exported to cryoSPARC; particles were re-extracted at a larger box size (box size 686 pixels, downsampled to 440 pixels) and *ab-initio* reconstructions followed by homogeneous refinements were used to verify assignments. Maps focused on leading ribosomes with E/E, P/P, and an empty A-site (+ effector) and E/E, P/P, A/A (-effector) were locally refined and combined as described above (**Figure S6F, S12**).

### Sample preparation for RF2 analysis by mass spectrometry

60 pmol purified RF2 (from the ΔRF1/2/3 PURExpress *in vitro* transcription/translation protein synthesis system [NEB]) were combined with 0.1 µM purified effector and 1 mM NAD^+^ in buffer H (50 mM HEPES-KOH pH 7.6, 10% glycerol, 5 mM Mg(OAc)_2_, 1 mM EDTA, 1 mM EGTA, 250 mM KCl) and incubated for 2 hours at 37 °C. Tryptic peptides were generated, reduced, alkylated, and desalted, starting from precipitated protein as performed previously^64^.

### LC–MS/MS of *in vitro* RF2 ADP-ribosylation reaction

Dried peptides (~2 mg) were resuspended in loading buffer (0.1% formic acid (FA) 4% Acetonitrile (ACN); 7 uL) before injection of 2 µL using an UltiMate 3000 RSLCnano system coupled to an Orbitrap Exploris 480 mass spectrometer (Thermo Fisher Scientific). Peptides were loaded and washed on a C18 trap column (Thermo Fisher Scientific, #164946) at 5 µL/min for 5 min in 96% buffer A (water, 0.1% Fa) and 4% buffer B (ACN, 0.1% FA). They were then resolved on a C18 analytical column (75 μm inner diameter × 50 cm, 2 μm particle size, 100 Å pore size; Thermo Fisher Scientific, #ES903) at 40 °C and 300 nL/min with a linear gradient of 4–21% B over 60 min, then 21–30% B over 10 min. MS1 scans (m/z 400–1200) were acquired at 120,000 resolution (at m/z 200) with an AGC target of 1 × 10^6^ and a maximum injection time of 50 ms. Data-dependent MS/MS used a 2-s cycle, targeting precursors of charge 2–6 or undetermined charge and greater than 5 × 10^4^ intensity, with automatic dynamic exclusion. Selected precursors were isolated with a 2-m/z window and fragmented by HCD at 26% normalized collision energy. Fragment spectra were acquired at 120,000 resolution (at m/z 200) with an AGC target of 1 × 10^5^ and an automatic maximum injection time (247 ms). All spectra were recorded in profile mode.

### ADP-ribosylation site identification

Mass-corrected MS2 spectra from the effector-treated and RF2-only DDA injections were searched for seven ADP-ribose marker ions (adenine, adenosine−H_2_O, adenosine, AMP, ADP, ADPr−2H_2_O and ADPr−H_2_O) within ±10 ppm of the expected m/z. This identified 8 spectra-precursor pairs with ≥3 markers in the effector-treated sample and none in the RF2-only sample. Each such precursor was mass-matched against tryptic peptides of the *E. coli* K-12 proteome (UniProt UP000000625) and RcaT-Eco9, plus reversed-sequence decoys, allowing ≤1 missed cleavage and fixed carbamidomethyl-Cys. This identified candidate peptides whose mass plus ADP-ribose (+541.0611 Da) was within ±10 ppm of the precursor. Candidates were scored by the number of backbone cleavages explained by b/y ions within ±10 ppm, with fragments unmodified or carrying ADP-ribose or any of its fragments. An assignment required outscoring all other targets, all reversed decoys and 50 sequence-shuffled decoys (p ≤ 0.02). One peptide, RF2 ^246^TSGAGGQHVNRTESAVR^262^ + ADP-ribose, met these criteria, in three spectra (scans 6185 and 6240, 3+; scan 6194, 4+). The highest signal spectrum (scan 6185) was then annotated for b/y ions carrying intact ADP-ribose or its fragments (ADPr−adenine, pyrophosphoribose, phosphoribose, phosphoribose−H_2_O, ribose), which localized the modification to His253. To test whether the modification was specific to the effector-treated sample, we extracted MS1 ion chromatograms of the modified precursor ([M+3H]^3+^ m/z 756.6445; [M+4H]^4+^ m/z 567.7352; ±10 ppm) from both injections. The precursor was detected only in the effector-treated sample, whereas the unmodified peptide was present in both.

## DATA AND CODE AVAILABILITY

Sequencing data are available in the Sequence Read Archive under BioProject PRJNA1538025. Summaries of spectral read counts and raw data for IP-MS/MS experiments and *in vitro* ADP-ribosylation of RF2 were deposited under MassIVE and can be accessed under accession MSV000103660. Density maps will be available through the EMDB as follows: 70S ribosome in the classical state (P/P, E/E) with and without nascent chain density (EMD-90115 and EMD-90109, respectively), 50S aberrant termination complexes 1-3 (EMD-90110, EMD-90111, and EMD-90112, respectively), and 30S pre-initiation complexes 1-2 (EMD-90114 and EMD-90113, respectively). Local refinements for leading and queuing ribosomes in disomes for effector-producing cells (EMD-90122 and EMD-90117, respectively), for IVTT reactions + effector (EMD-90126 and EMD-90124, respectively), and for IVTT reactions − effector (EMD-90125 and EMD-90123, respectively). All other data are available in the manuscript or supplementary materials. This paper does not report any original code.

## AUTHOR CONTRIBUTIONS

All experiments were conceived and designed by C.R.B and M.T.L, and performed by C.R.B., except for cryoEM and mass spectrometry analysis of RF2, which were done by M.B.M. and J.H.D. Figure design, manuscript writing, and editing done by C.R.B., M.B.M., J.H.D, and M.T.L. Overall project supervision and funding provided by M.T.L. with contributions from J.H.D.

## ACKNOWLEDGMENTS

The authors thank Gene-Wei Li, Jay Hesselberth, Allen Buskirk, Ian Roney, and all Laub lab members for valuable discussions; Steve Bell and Shiva Dahagam for assistance with yeast cloning and protein purification; and Tong Zhang and Miles Black for critical readings of the manuscript. Plasmids pTU175 and pJB145 were a gift from Jacob Bobonis and Athanasios Typas. The authors used AI-assisted tools (OpenAI, Anthropic) for some scripting in figure construction and cryoEM data analysis. Manuscript was typeset using mudbrick (https://github.com/Under-Refinement/mudbrick), developed by Robert C. Miller.

