## Supplemental Figures and Tables for "Retron-Eco9 ADP-ribosylates release factors to disrupt translation during phage infection"

### SUPPLEMENTARY FIGURES

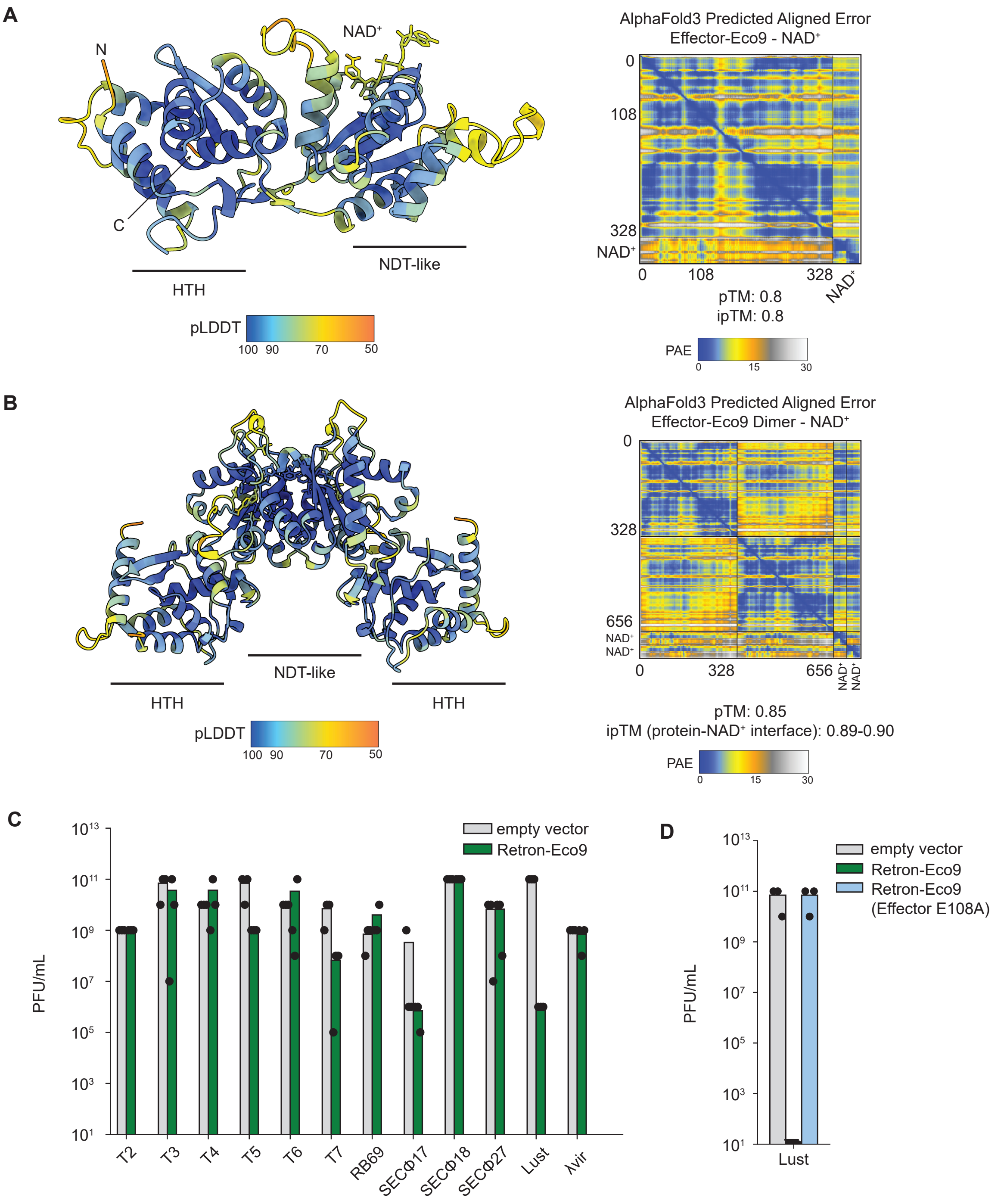


**Figure S1. Structural predictions of Effector-Eco9. (A)** Left, AlphaFold3 prediction of Effector-Eco9-NAD⁺ colored by pLDDT confidence. Right, PAE plot for predicted complex. **(B)** Left, AF3 prediction of Effector-Eco9-NAD⁺ dimer colored by pLDDT confidence. Right, PAE plot for predicted structure. **(C)** Summary of three independent replicates of phage plaquing assays in Figure 1B. **(D)** Summary of three independent replicates of phage plaquing assays in Figure 1C.

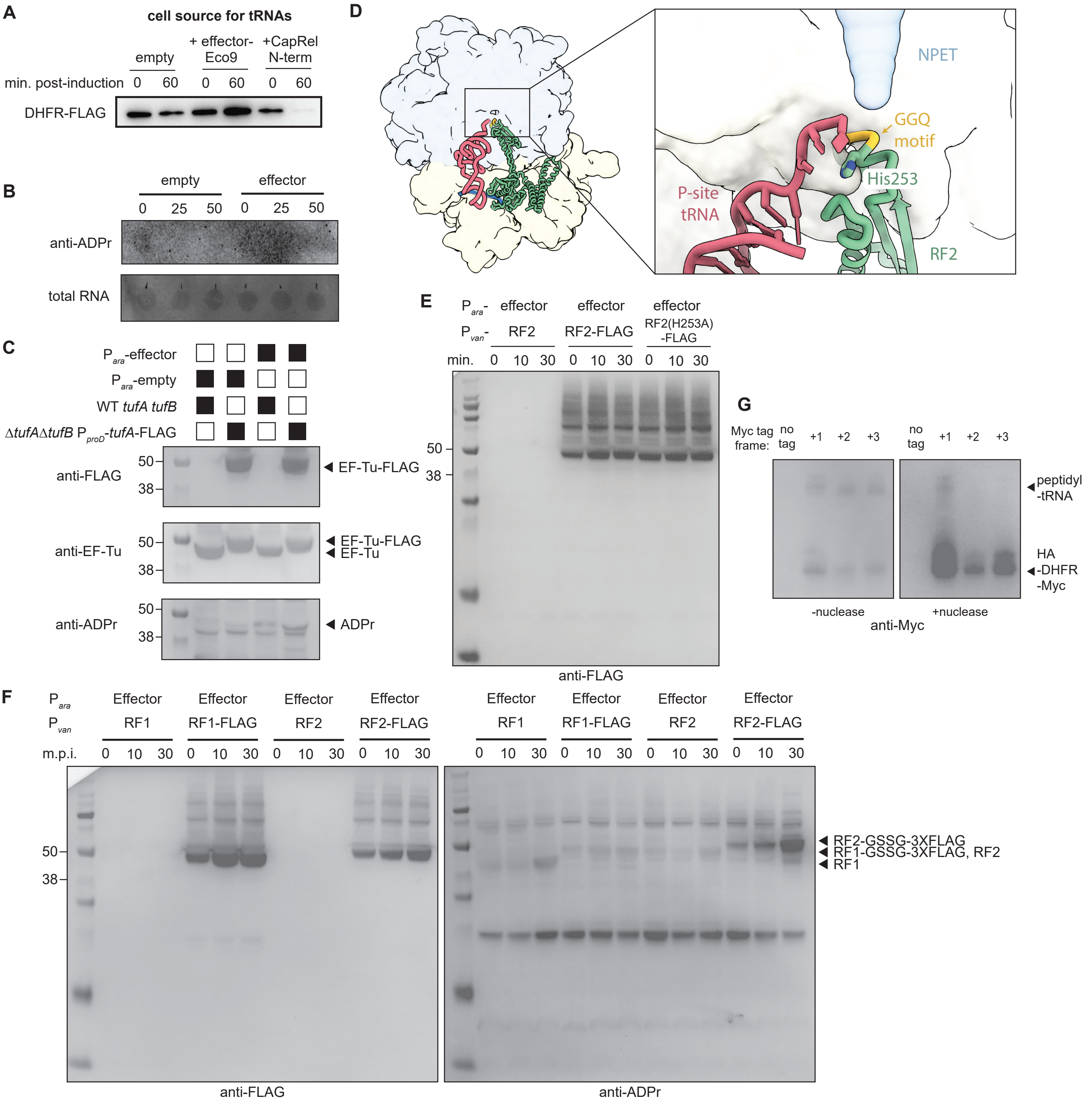


**Figure S2. Analysis of Effector-Eco9 ADP-ribosyltransferase activity. (A)** Anti-FLAG immunoblot assessing production of DHFR-FLAG in an IVTT reaction. IVTT exclusively used tRNA harvested from cells producing effector-Eco9 or the CapRel$^{SJ46}$ N-term from an arabinose-inducible promoter at the times indicated post-addition of arabinose. Representative of two biological replicates. **(B)** Top, RNA anti-ADPr immunoblot with total RNA extracted from cells expressing Effector-Eco9 from an arabinose-inducible promoter at the time indicated post-arabinose addition. Bottom, methylene blue stain for total RNA (enhanced to increase blot contrast). **(C)** Immunoblot to detect modification of EF-Tu. Cells containing wild-type or FLAG-tagged EF-Tu and expressing Effector-Eco9 from an arabinose-inducible promoter were harvested 30 minutes post-arabinose addition. Cell lysates were blotted for anti-FLAG, anti-EF-Tu, and anti-ADPr. **(D)** Atomic model (PDB: 6OUO) of 70S ribosome bound to RF2 (green), with $^{250}$GGQ$^{252}$ colored gold, P-site tRNA colored red, and H253 shown in stick representation. Exit tunnel is highlighted with a blue semi-transparent mask, and large and small subunits colored blue and yellow, respectively. **(E)** anti-FLAG immunoblot corresponding to samples in Figure 2F. **(F)** Immunoblot to detect ADP-ribosylation of RF1 and RF2. Cells expressing wild-type or FLAG-tagged RF1 or RF2 from a vanillic acid-inducible promoter and Effector-Eco9 from an arabinose-inducible promoter were harvested at the times indicated post-arabinose addition. Cell lysates were blotted for anti-FLAG and anti-ADPr. **(G)** Immunoblot to detect HA-DHFR-Myc from *in vitro* transcription/translation reactions lacking RF1 and RF2, corresponding to Figure 3C. Following the reaction, samples were treated with nuclease and then blotted again for anti-Myc signal.

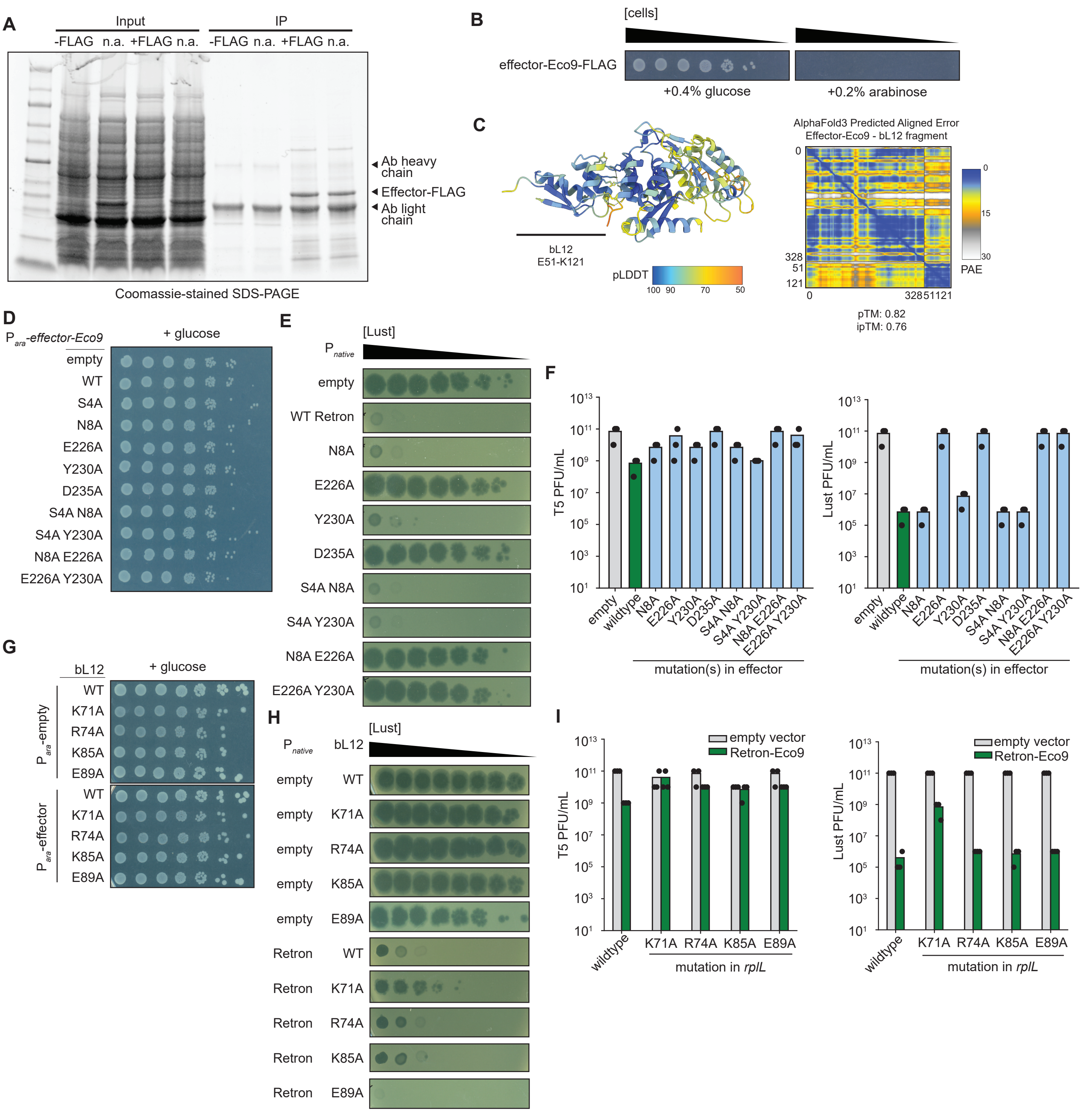


**Figure S3. Analysis of interactions between Effector-Eco9 and bL7/bL12. (A)** Coomassie-stained gel of cell lysates (i.e. input) and immunoprecipitated (i.e. IP) fractions from cells expressing Effector-Eco9(E108A) or Effector-Eco9(E108A)-FLAG from an arabinose-inducible promoter 140 minutes post-addition of arabinose. **(B)** Cell viability as assessed by serial dilution for cells expressing Effector-Eco9-FLAG from an arabinose-inducible promoter. **(C)** AF3-predicted structure for Effector-Eco9 and C-term fragment (residues 51-121) of bL7/bL12 dimer colored by pLDDT confidence and PAE plot for predicted bL7/bL12 dimer fragment-Effector-Eco9 structure. **(D)** Cell viability as assessed by serial dilution for cells expressing wild-type or mutant versions of Effector-Eco9 from an arabinose-inducible promoter in glucose-containing media. Corresponds to panel Figure 4D. **(E)** Serial dilutions of Lust phage spotted on lawns of cells containing plasmids expressing wild-type Retron-Eco9 or Retron-Eco9 containing the indicated mutations in Effector-Eco9. **(F)** Summary of three independent replicates of phage plaquing assays in Figure 4E (right) and Figure S4E (left). **(G)** Cell viability assessed by serial dilutions for strains containing indicated mutations in genomic *rplL* locus (encoding bL7/bL12) and plasmids expressing Effector-Eco9 from an arabinose-inducible promoter or an empty vector in glucose-containing media. Corresponds to panel Figure 4F. **(H)** Serial dilutions of Lust phage spotted on lawns of cells containing indicated mutations in *rplL* locus and plasmids expressing wild-type Retron-Eco9 or an empty vector. **(I)** Summary of three independent replicates of phage plaquing assays in Figure 4G (left) and Figure S4H (right).

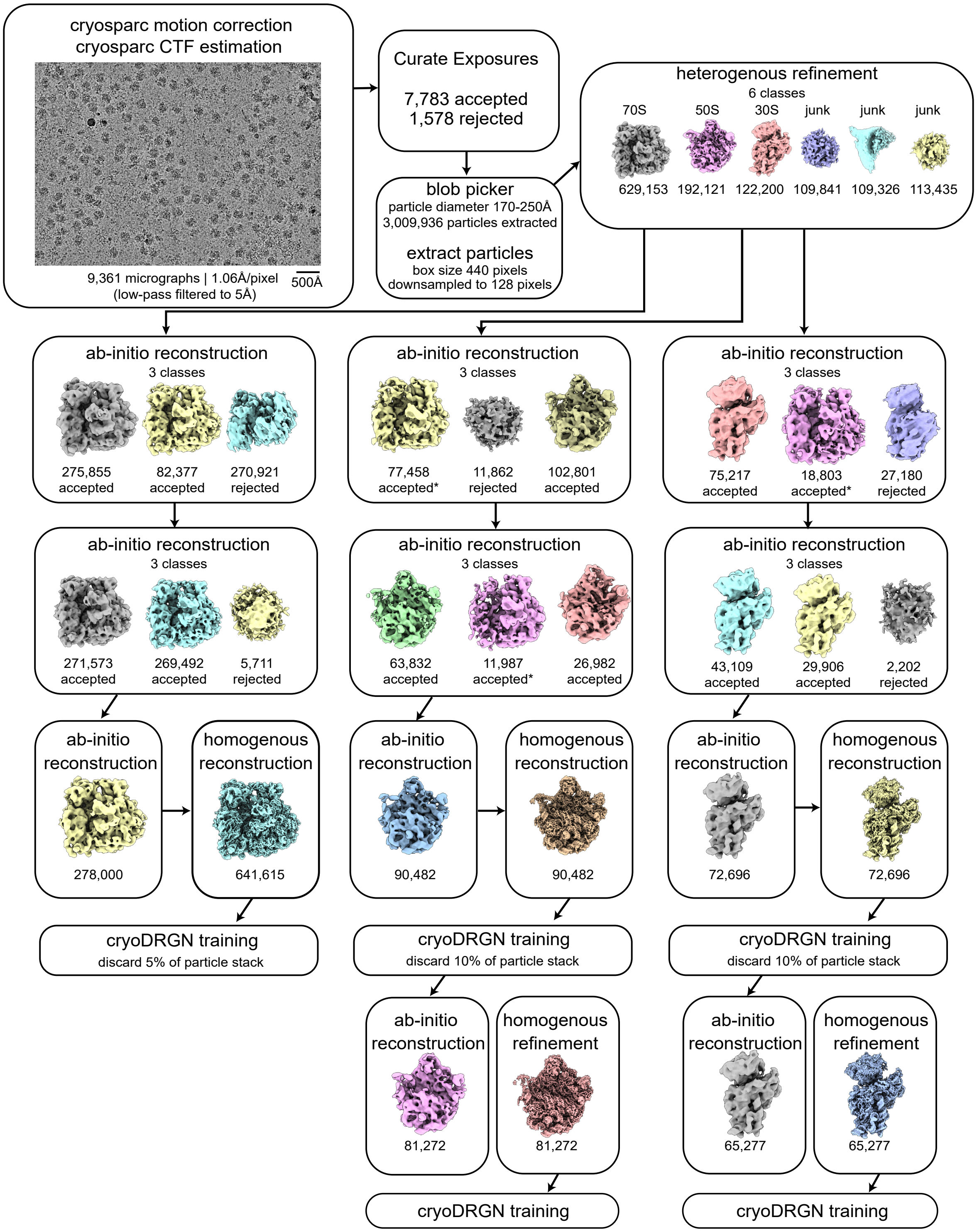


**Figure S4. CryoSPARC pre-processing workflow for analysis of +effector cryoPRISM dataset.** The job names and key parameters are listed, with particle numbers listed and representative maps depicted. Accepted particles with * indicates mis-classified 70S particles, which were subsequently transferred into the 70S classification workflow.

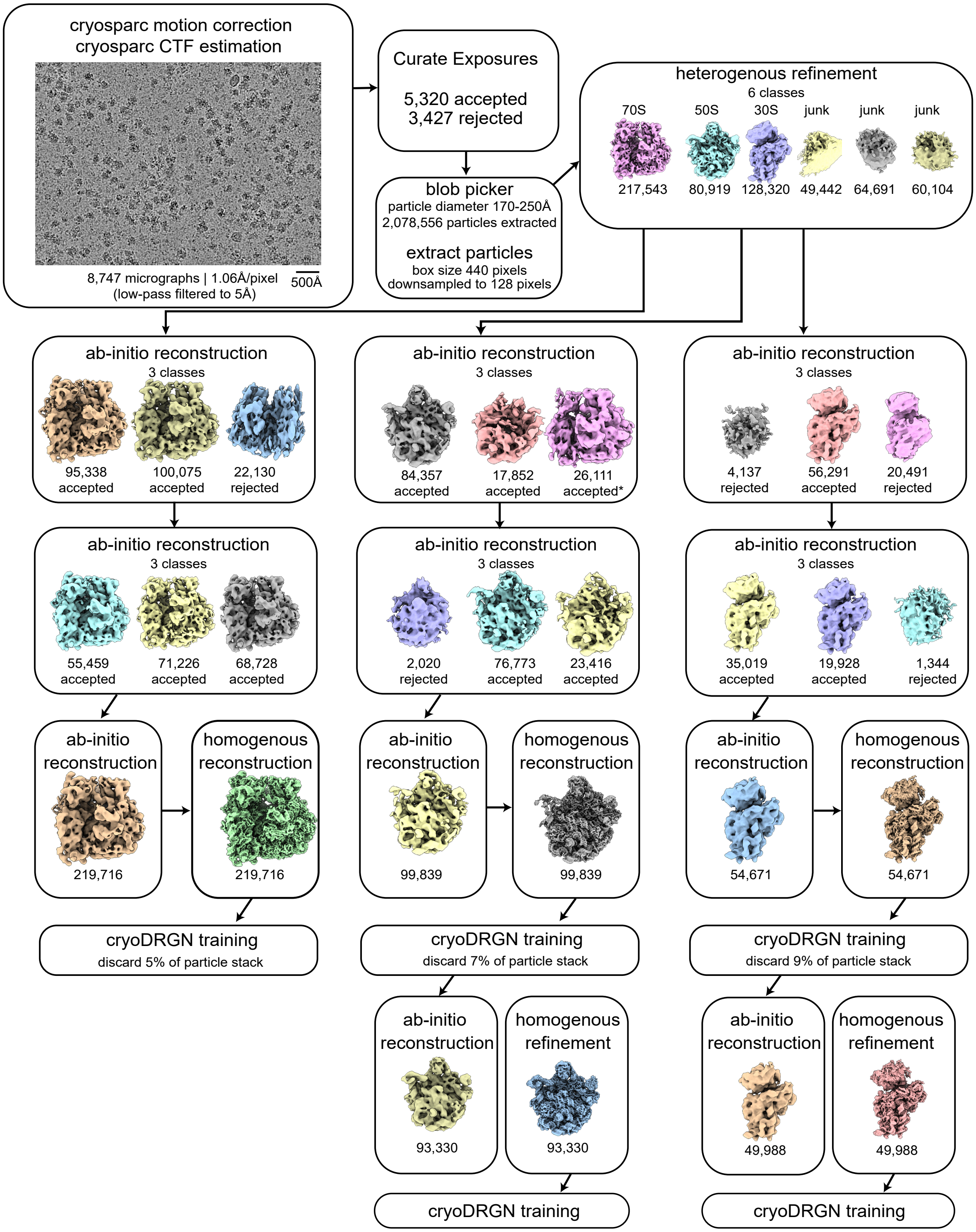


**Figure S5. CryoSPARC pre-processing workflow for analysis of -effector cryoPRISM dataset.** The job names and key parameters are listed, with particle numbers listed and representative maps depicted. Accepted particles with * indicates mis-classified 70S particles, which were subsequently transferred into the 70S classification workflow.

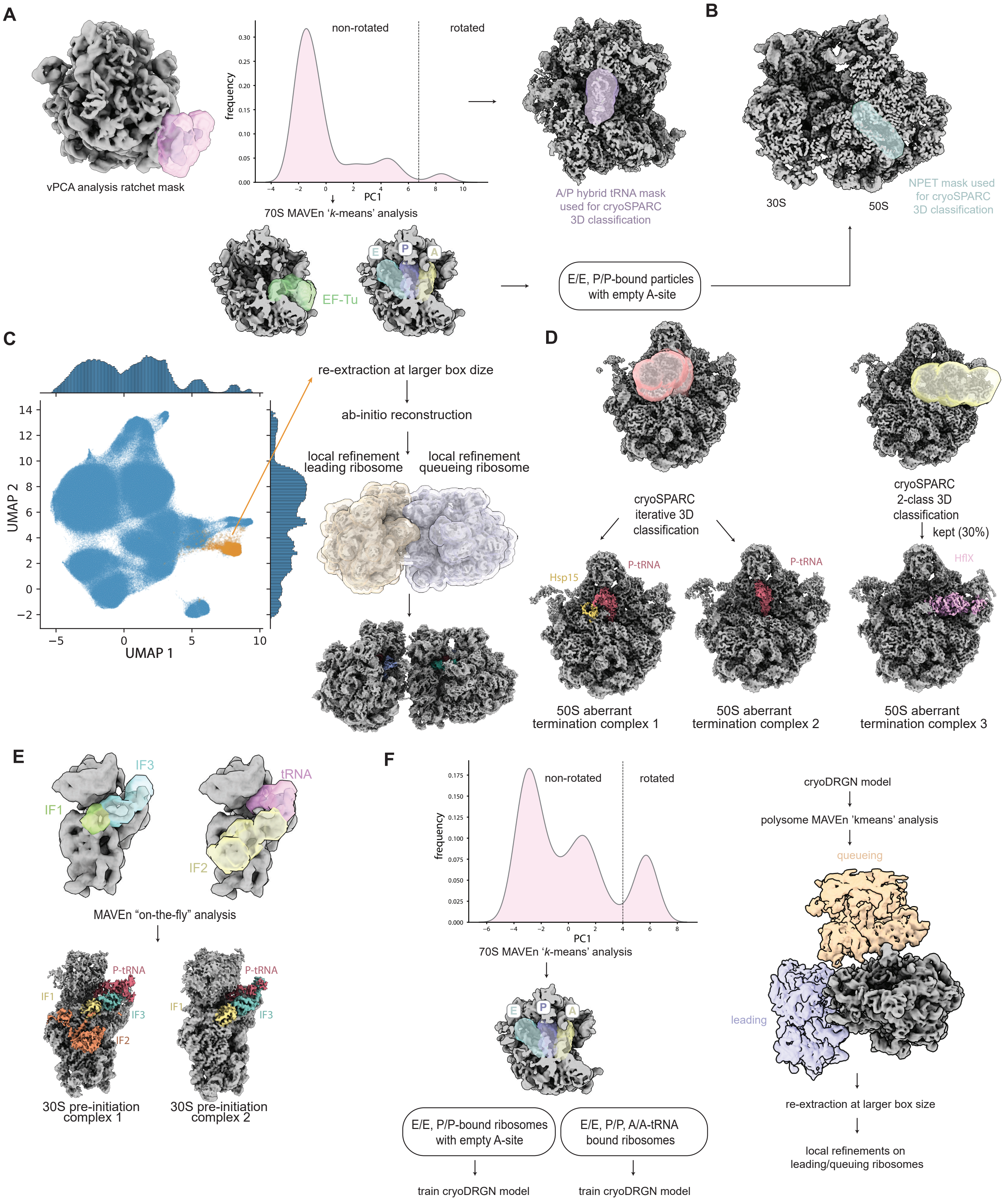


**Figure S6. Image processing workflow for analysis of 70S, 50S, 30S, and disome/polysome structural states. (A)** Voxel PCA analysis (vPCA) was employed to capture the transition of the small subunit relative to the large subunit during ribosome ratcheting using voxels within a mask encompassing 16S rRNA helices 6, 10, and 17 (PDBs 7ST6 and 7SSN), and the resulting particle density along the first principal component is plotted. A threshold used to separate particles in the rotated conformation bearing tRNAs in the hybrid state is drawn. Subsequent classification of A/P site-tRNAs was performed in cryoSPARC using the mask shown. (*overleaf*)

**Figure S6** ***(continued from prior page).*** For 'non-rotated' particles below the threshold, masks utilized for MAVEn comprising E/E, P/P, and A/A-site tRNAs in the classical conformation and EF-Tu are shown, with occupancy thresholds given in SI Tables 2,3, and 6. **(B)** Mask spanning nascent chain exit tunnel (NPET) was generated based on a poly-alanine backbone built into exit tunnel and used to classify E/E, P/P-tRNA-bound 70S particles via cryoSPARC's 3D classification job. **(C)** Scatter plot with marginal distributions of cryoDRGN latent space for all 70S ribosomes after UMAP dimensionality reduction, where each point marks a projected particle from the (+) effector cryoPRISM dataset. Disome particles (colored in orange) from Figure 6E were identified by inspection of 50 *k*-means centroid volumes and used for subsequent cryoSPARC re-extraction, ab-initio reconstruction, and focused refinement. **(D)** Masks spanning the binding sites for Hsp-15/P-site tRNA (left) and the PTC (right) were used in cryoSPARC 3D classification jobs to curate particles stacks for 50S aberrant termination complexes 1-3 (Fig 6C). **(E)** Masks used for MAVEn "on-the-fly" analysis of 30S particles encompassing IF1, IF2, IF3, and initiator tRNA to classify particles into 30S pre-initiation complexes 1 and 2 as well as 30S subunits bound by IF1/IF3 and without IFs bound (Figure 6A). **(F)** Workflow for IVTT 70S particle classification. Particles were separated based on the 30S rotational state relative to the 50S using vPCA analysis with 1,000 *k*-means volumes and subsequently classified for A/A and P/P tRNA occupancy using MAVEn "k-means" analysis. Separate cryoDRGN models were trained on ribosomes with and without A-site tRNA occupancy and used to generate another 1,000 *k*-means volumes for each sub-class. Masks used for MAVEn "*k*-means" analysis of disomes on IVTT datasets are shown. Particles with adjacent density for a leading ribosome, queueing ribosome, or both were re-extracted at a larger box size in cryoSPARC and locally refined on each monosome.

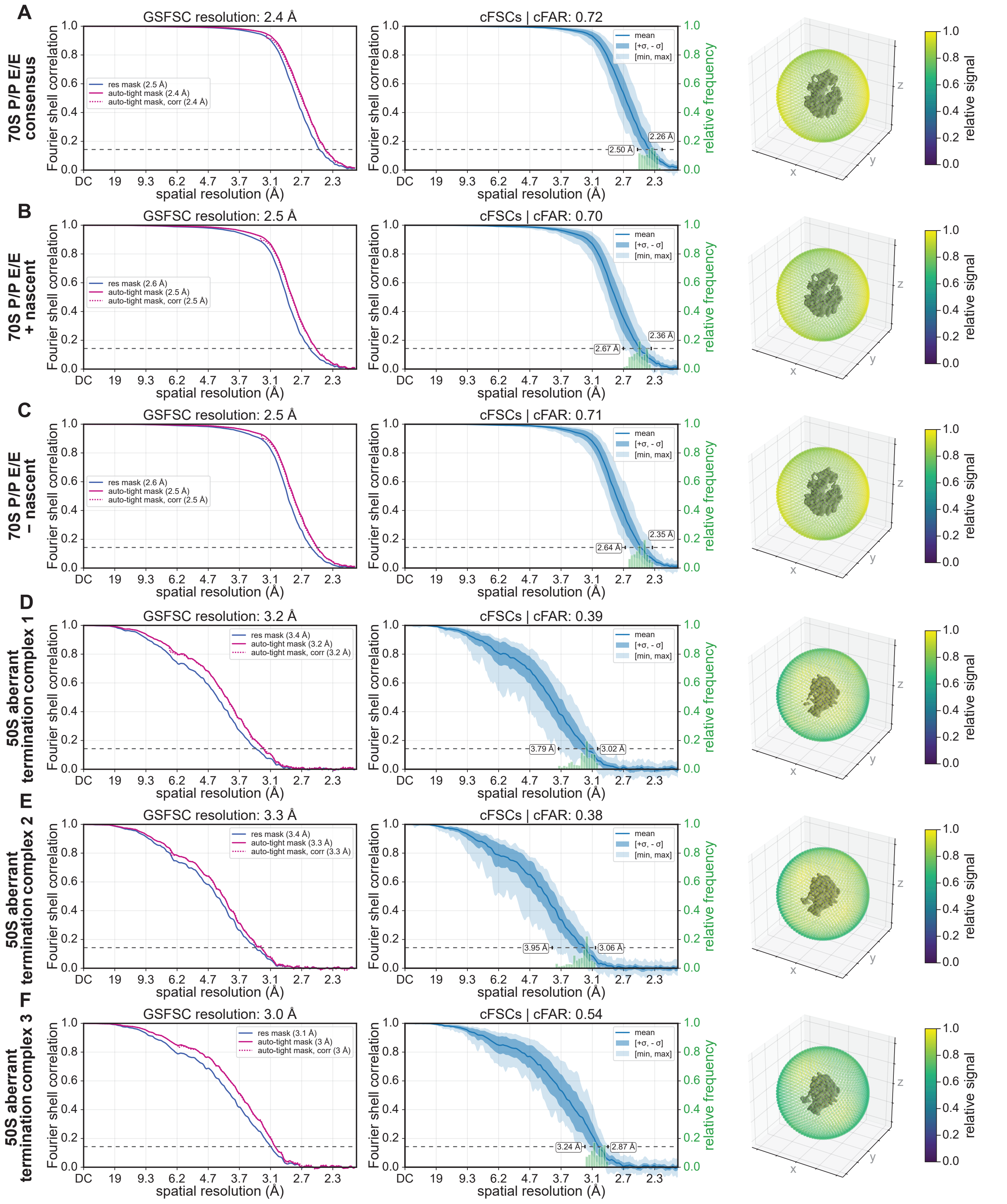


**Figure S7. Quality assessment of consensus 3D reconstructions of 70S and 50S states from cryoPRISM +effector dataset.** Global resolution estimates calculated using Fourier shell correlation (FSC) between independently refined half-maps as implemented in cryoSPARC's homogeneous refinement method. The FSC = 0.143 threshold is indicated with a dashed line, and the corresponding resolution is reported (left). Conical FSC (cFSC) curves (mean, min, max, ±1σ, as listed in legend) are plotted, with the cFSC area ratio (cFAR) shown in the title and the directional resolution range labeled on the figure. The relative frequency of cFSC values across individual conical slices is shown as a histogram (middle). A sphere colored by relative signal is visualized surrounding each density map (right). Plots are shown for 70S (A-C) and 50S states (D-F) depicted in Figure 6B-C.

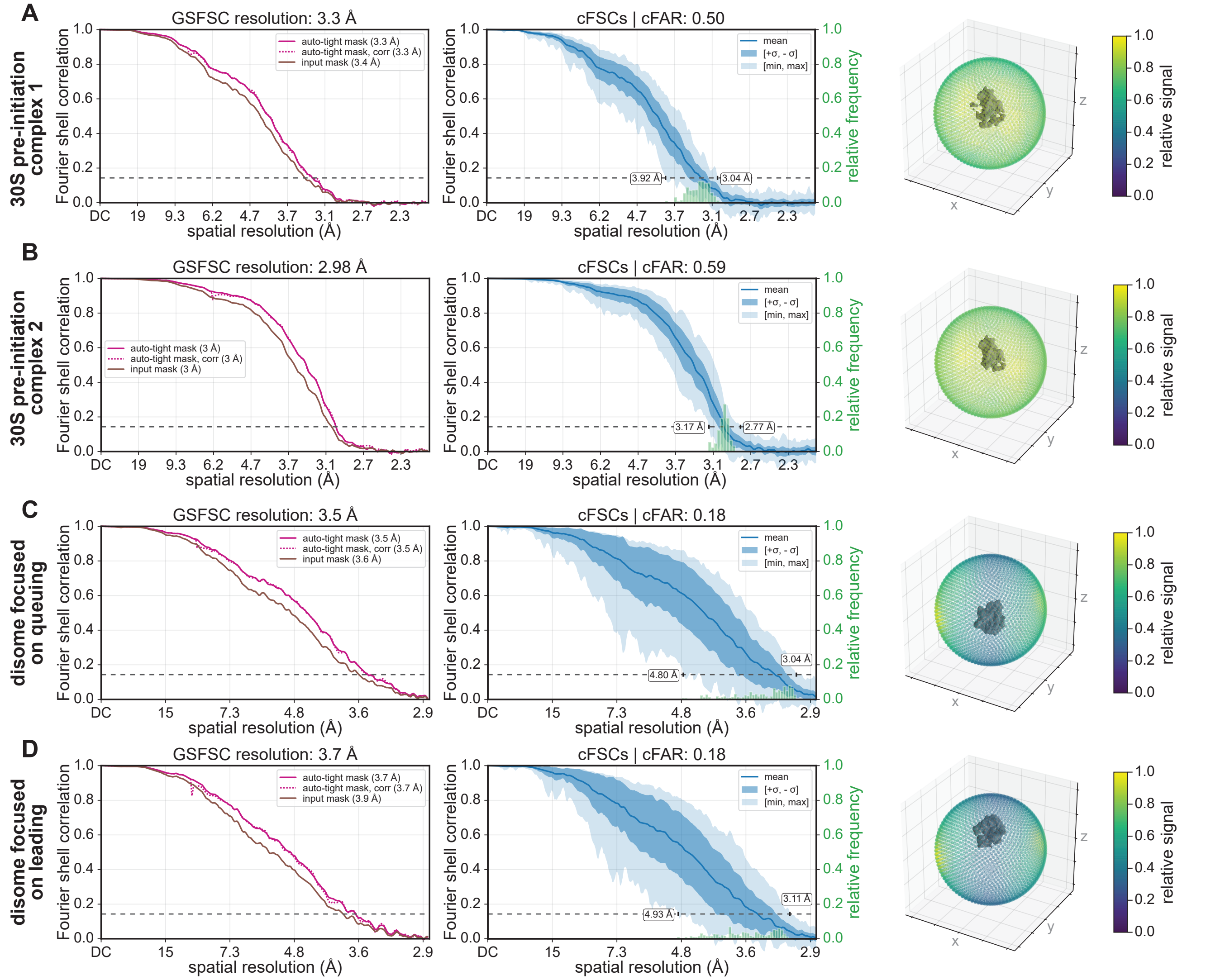


**Figure S8. Quality assessment of consensus 3D reconstructions of disomes and 30S states from cryoPRISM +effector dataset.** Global resolution estimates calculated using Fourier shell correlation (FSC) between independently refined half-maps as implemented in cryoSPARC's homogeneous refinement method. The FSC = 0.143 threshold is indicated with a dashed line, and the corresponding resolution is reported (left). Conical FSC (cFSC) curves (mean, min, max, ±1σ, as listed in legend) are plotted, with the cFSC area ratio (cFAR) shown in the title and the directional resolution range labeled on the figure. The relative frequency of cFSC values across individual conical slices is shown as a histogram (middle). A sphere colored by relative signal is visualized surrounding each density map (right). Plots are shown for 30S pre-initiation complexes 1-2 (A-B) and disomes (C-D) depicted in Figure 6D-E.

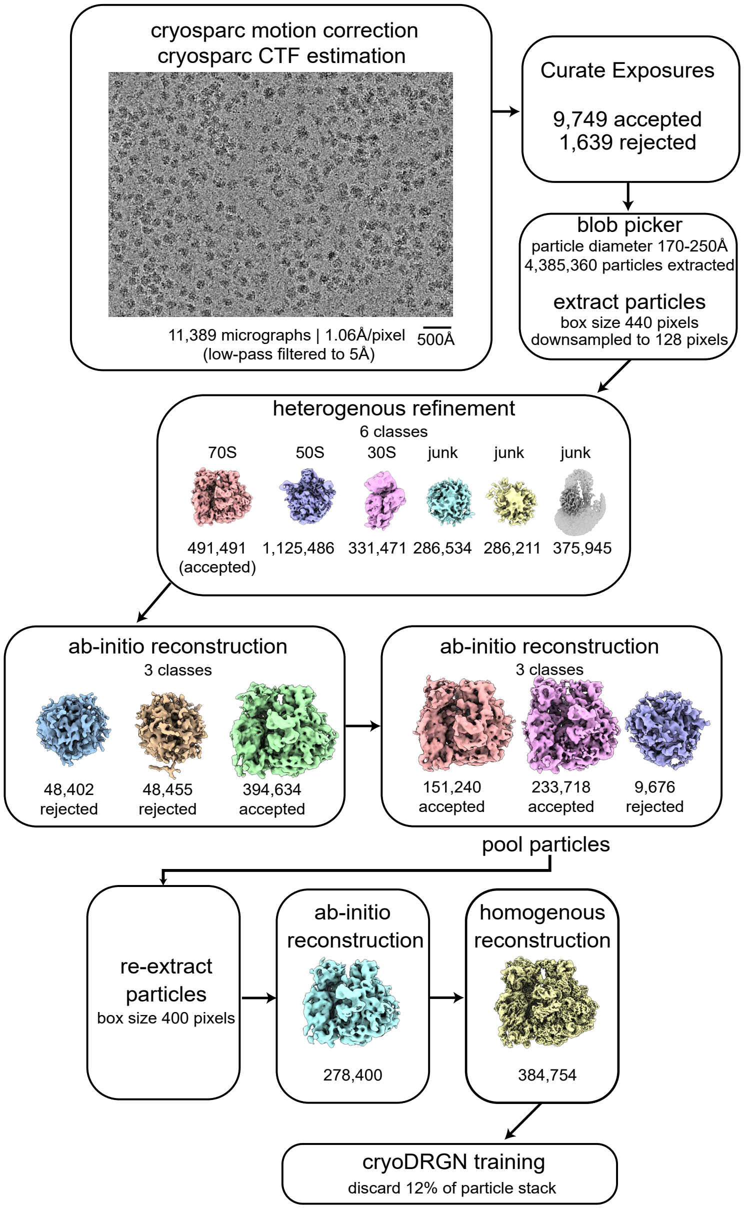


**Figure S9. CryoSPARC pre-processing workflow for analysis of IVTT +effector dataset.** The job names and key parameters are listed, with particle numbers listed and representative maps depicted.

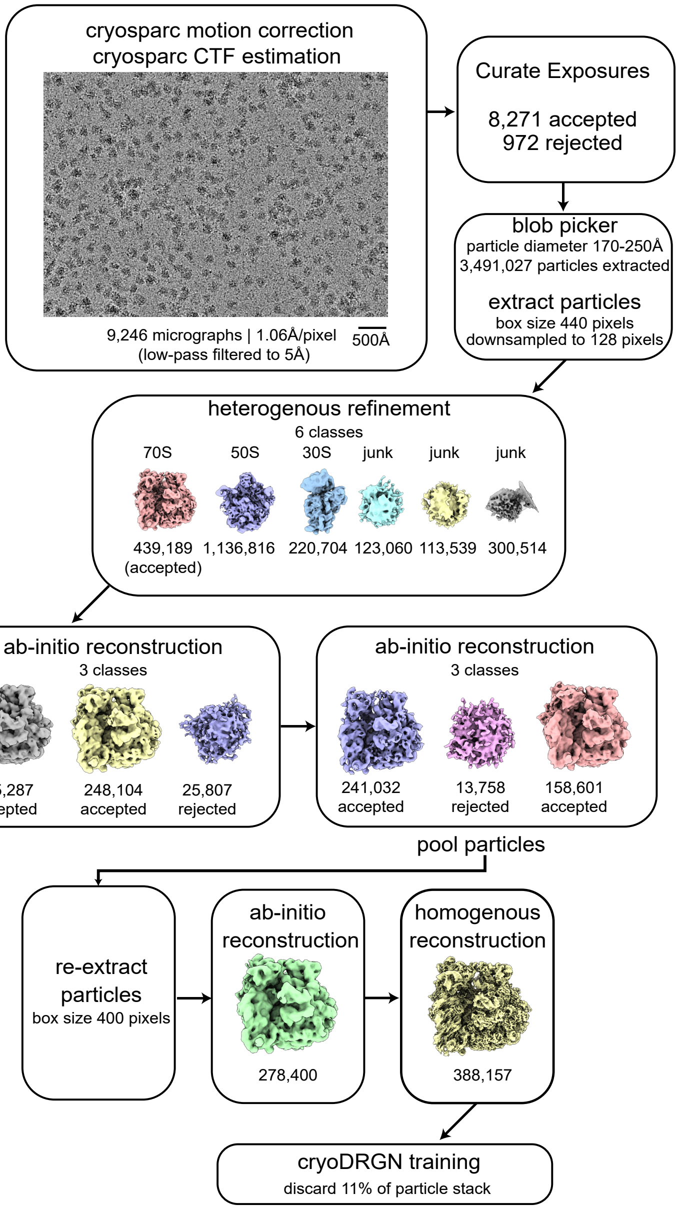


**Figure S10. CryoSPARC pre-processing workflow for analysis of IVTT -effector dataset.** The job names and key parameters are listed, with particle numbers listed and representative maps depicted.

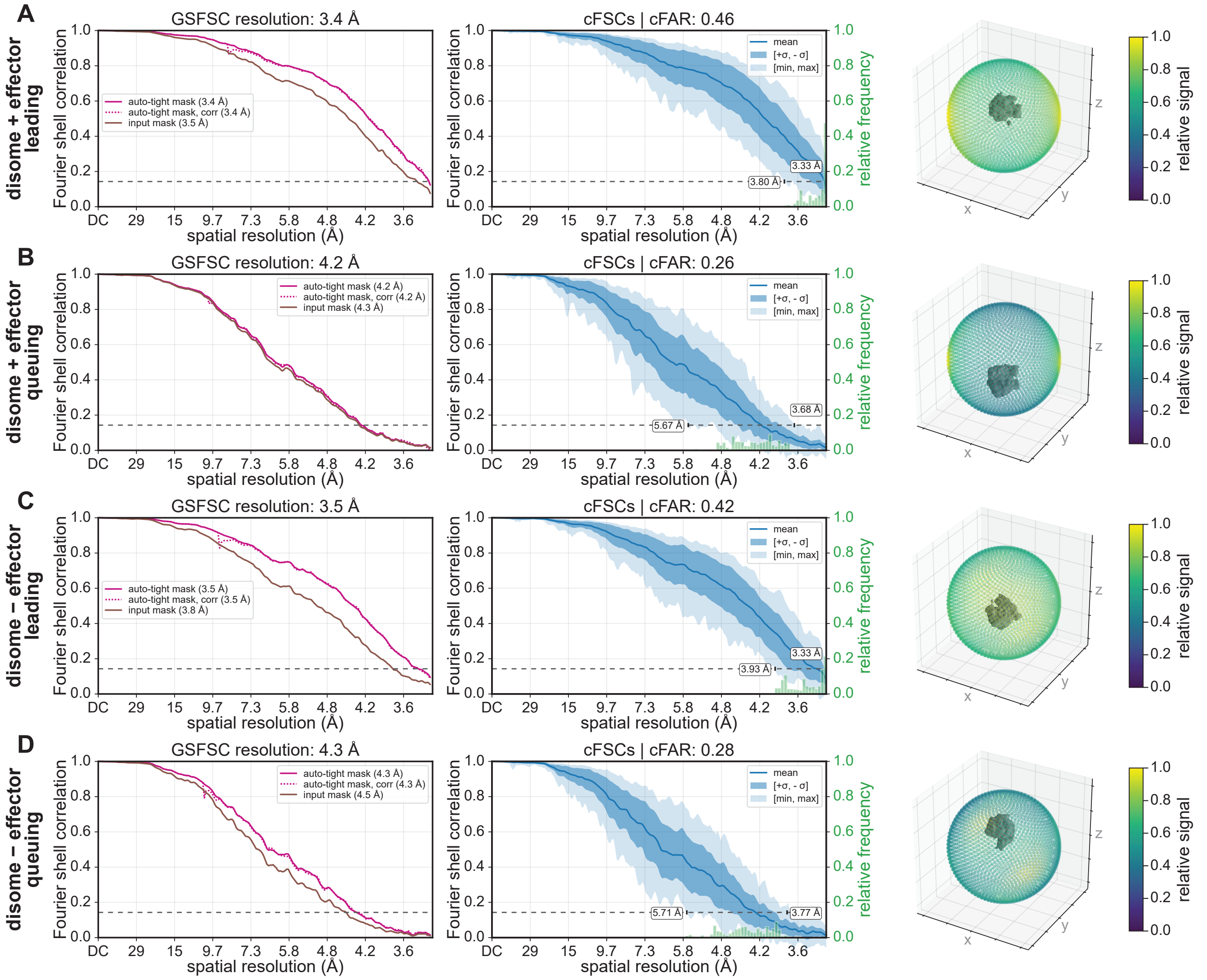


**Figure S11. Quality assessment of consensus 3D reconstructions of disomes from IVTT datasets.** Global resolution estimates calculated using Fourier shell correlation (FSC) between independently refined half-maps as implemented in cryoSPARC's homogeneous refinement method. The FSC = 0.143 threshold is indicated with a dashed line, and the corresponding resolution is reported (left). Conical FSC (cFSC) curves (mean, min, max, ±1σ, as listed in legend) are plotted, with the cFSC area ratio (cFAR) shown in the title and the directional resolution range labeled on the figure. The relative frequency of cFSC values across individual conical slices is shown as a histogram (middle). A sphere colored by relative signal is visualized surrounding each density map (right). Plots are shown for disomes from the + effector (A-B) and – effector (C-D) IVTT cryoEM maps in Figure S12.

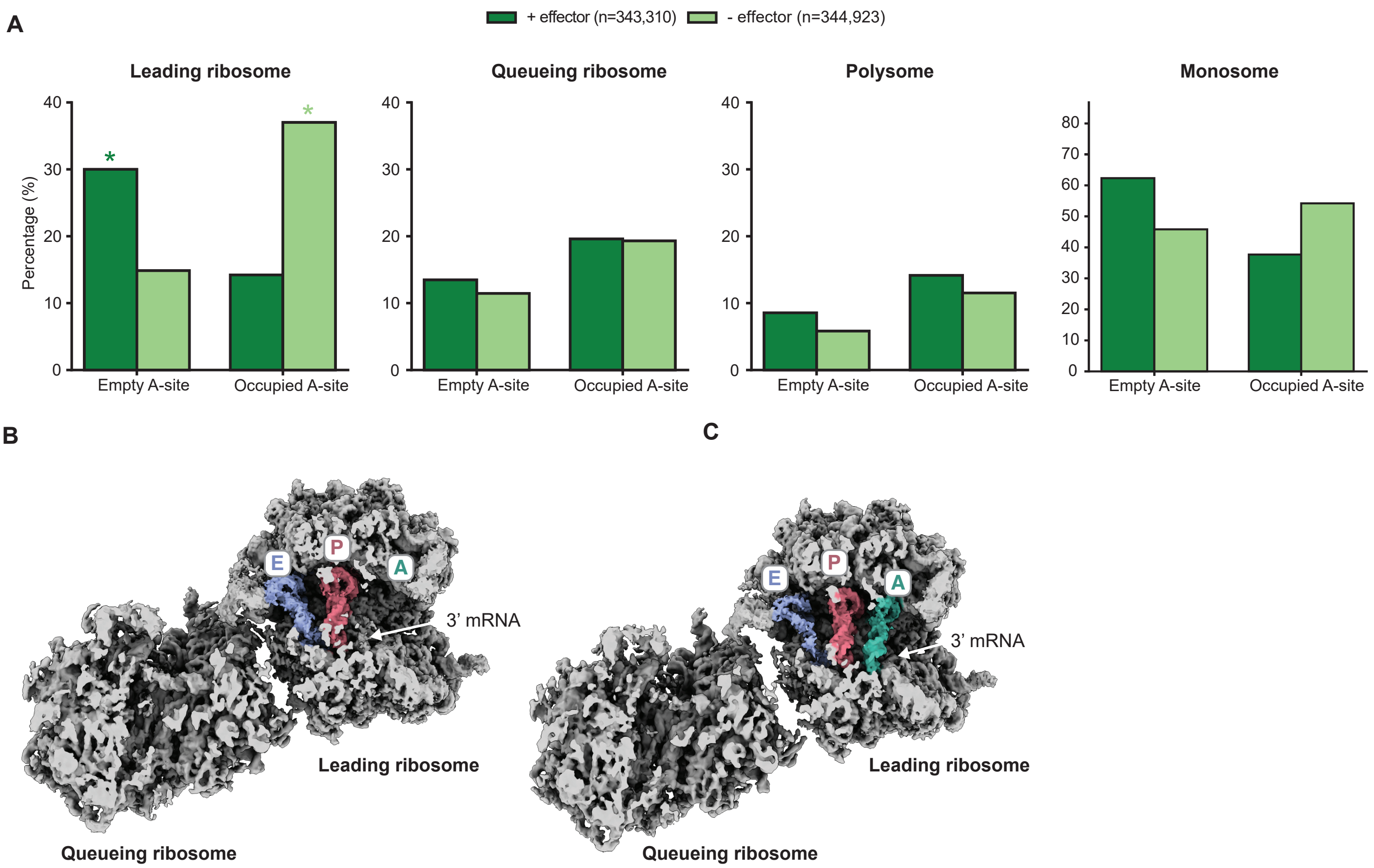


**Figure S12. CryoEM of IVTT reactions with purified effector indicates enrichment of leading ribosomes on mRNA with empty A-site.** **(A)** Bar chart depicting frequency of particles that are members of monosome, disomes, or polysomes with and without A-site occupancy, identified by cryoEM from IVTT reaction. IVTT reactions are supplemented with HA-DHFR-TGA template, $NAD^+$, and purified effector (+ effector, dark green) or without effector (- effector, light green). Frequencies of particles with and without A-site tRNA that are identified as the leading member of a disome pair, queueing member of a disome pair, polysomes (neighboring ribosomes observed in leading and queueing position), or monosomes are plotted. Percentages of each state computed relative to the sum of disomes/polysomes. Dark and light green asterixis indicate cryoEM reconstructions shown in (B) and (C), respectively. **(B)** Density map for the most abundant disome state from the + effector condition, shown in top-down clipped view. P and E-site tRNAs, the empty A-tRNA binding site, and the 3' end of the mRNA are labeled and colored as in Fig 6. **(C)** Density map for the most abundant disome state from the - effector condition, shown in top-down clipped view. A, P, E-site tRNAs and the 3' end of the mRNA are labeled and colored as in (C).

### SUPPLEMENTARY TABLES

| 70S particle origin | $1^{st}$ principal component threshold for vPCA |
|---|---|
| + effector cryoPRISM | 6.75 |
| - effector cryoPRISM | 5.9 |
| + effector IVTT | 4 |
| - effector IVTT | 3.6 |

**Table S1.** Thresholds used to distinguish 30S rotated state (with hybrid tRNAs) relative to the 50S.

| | tRNA | | translation factors |
|---|---|---|---|
| State | P-site (PDB: 7st6) | A-site (PDB: 7st6) | EF-Tu PDB: 6wd2 |
| EF-Tu, P/P, E/E | * | < 0.25 | > 0.1 |
| A/A, P/P, E/E | * | > 0.25 | < 0.25 |
| P/P, E/E | * | < 0.25 | > 0.1 |

**Table S2.** Thresholds used to define 70S populations (+ effector). *indicates no threshold is set due to uniform presence of feature.

| | tRNA | | translation factors |
|---|---|---|---|
| State | P-site (PDB: 7st6) | A-site (PDB: 7st6) | EF-Tu PDB: 6wd2 |
| EF-Tu, P/P, E/E | > 0.25 | < 0.25 | > 0.2 |
| A/A, P/P, E/E | > 0.25 | > 0.25 | < 0.2 |
| P/P, E/E | > 0.25 | < 0.25 | < 0.2 |
| idle | < 0.25 | | |

**Table S3.** Thresholds used to define 70S populations (- effector).

| | tRNA | initiation factors | | |
|---|---|---|---|---|
| State | Initiator P-site | IF1 PDB: 5lmp | IF2 PDB: 5lmp | IF3 PDB: 5lmp |
| PIC complex 1 | | | > 0.35 | |
| PIC complex 2 | > 0.25 | > 0.25 | < 0.35 | > 0.3 |
| IF1 and IF3 | < 0.25 | > 0.25 | < 0.35 | > 0.3 |
| unbound | | | remaining particles | |

**Table S4.** Thresholds used to define 30S populations (+ effector).

| | tRNA | initiation factors | | |
|---|---|---|---|---|
| State | Initiator tRNA | IF1 PDB: 5lmp | IF2 PDB: 5lmp | IF3 PDB: 5lmp |
| PIC complex 1 | | | > 0.4 | |
| PIC complex 2 | > 0.35 | > 0.45 | < 0.4 | > 0.35 |
| IF1 and IF3 | < 0.35 | > 0.45 | < 0.4 | > 0.35 |
| unbound | | | remaining particles | |

**Table S5.** Thresholds used to define 30S populations (- effector).

| | tRNA | | neighboring ribosome | |
|---|---|---|---|---|
| State | P-site (PDB: 7st6) | A-site (PDB: 7st6) | queueing | leading |
| P/P, E/E leading | * | < 0.25 | (+) | (-) |
| A/A, P/P, E/E leading | * | > 0.25 | (+) | (-) |
| P/P, E/E queueing | * | < 0.25 | (-) | (+) |
| A/A, P/P, E/E queueing | * | > 0.25 | (-) | (+) |
| A/A, P/P polysome | * | < 0.25 | (+) | (+) |
| A/A, P/P, E/E polysome | * | > 0.25 | (+) | (+) |

**Table S6.** Thresholds used to define IVTT 70S monosome, disome, and polysome populations (+ and - effector). *indicates no threshold is set due to uniform presence of feature. Note that same thresholds were used for both + effector and – effector IVTT datasets.
